# Transcription factor exchange enables prolonged transcriptional bursts

**DOI:** 10.1101/2023.05.15.540758

**Authors:** Wim Pomp, Joseph V.W. Meeussen, Tineke L. Lenstra

**Affiliations:** Division of Gene Regulation, The Netherlands Cancer Institute, Oncode Institute, Plesmanlaan 121, 1066CX Amsterdam, The Netherlands

**Keywords:** transcription factor, transcriptional bursting, single-molecule imaging, DNA binding, kinetics, cooperativity

## Abstract

Single-molecule imaging inside living cells has revealed that transcription factors (TFs) bind to DNA transiently, but a long-standing question is how this transient binding is related to transcription activation. Here, we devised a microscopy method to simultaneously measure transient TF binding at a single locus and the effect of these binding events on transcription. We show that DNA binding of the yeast TF Gal4 activates transcription of a target gene within a few seconds, with at least ^∼^20% efficiency and with a high initiation rate of ^∼^1 RNA/s. Gal4 DNA dissociation decreases transcription rapidly. Moreover, at a gene with multiple binding sites, individual Gal4 molecules only rarely stay bound throughout the entire burst, but instead frequently exchange during a burst to increase the transcriptional burst duration. Our results suggest a mechanism for enhancer regulation in more complex eukaryotes, where TF cooperativity and exchange enable robust transcription activation.

## Introduction

Gene-specific transcription factors (TFs) bind to specific DNA sequences in DNA regulatory regions of target genes to control gene expression. During their search for specific target sequences, TFs diffuse through the nucleoplasm and associate non-specifically with nucleosomes, DNA-bound proteins and non-target DNA sequences^**1–3**^. Upon binding to their target sequence, activating TFs recruit cofactors and the preinitiation complex to activate transcription. Single-cell and single-molecule imaging has revealed that these processes are highly dynamic inside living cells. The majority of TFs were found to bind to DNA for only a few seconds ^**4**^. Furthermore, for many genes, transcription is discontinuous and occurs in bursts ^**5**,**6**^. Transcription levels are determined by the frequency of the bursts, the duration of the bursts, and the number of polymerases that initiate during bursts (burst size). The bursting dynamics are to a large degree a consequence of the properties of the TFs and their interactions with the DNA, but the exact kinetic relationship is still unclear ^**7–10**^.

Even though TF binding and transcriptional bursting are both dynamic, they occur on different time scales: the duration of TF binding is on the order of seconds, while the duration of a transcriptional burst is on the order of minutes. A fundamental outstanding question is how these two timescales can be connected. In other words, how can such transient TF binding robustly activate transcription? In literature, three kinetic models have been proposed to bridge this timescale disconnect between TF binding and transcription ^11–13^ (Figure 1). In the first model, TFs bind throughout the entire burst, such that their residence time determines both the start and end of a burst. Although the majority of TFs bind DNA for a few seconds, the imaging methods used to determine these residence times are photobleaching prone and hence could have missed a subpopulation of TF binding events with longer residence times. In the second model, TF binding initiates a multi-step process with recruitment of other factors, which activate transcription even after the TF has left. TF binding would then be required to start a burst, but not to end It. In the third model, cooperative binding of TFs results in longer periods of high promoter occupancy, during which individual TFs have short residence times, but constantly exchange. Such cooperative binding may be facilitated by self-interactions or clustering of TFs near the target gene ^14,15^. Due to technical limitations, these kinetic models for transcription activation have so far only scarcely been tested ^16–18^.

**Figure 1.**
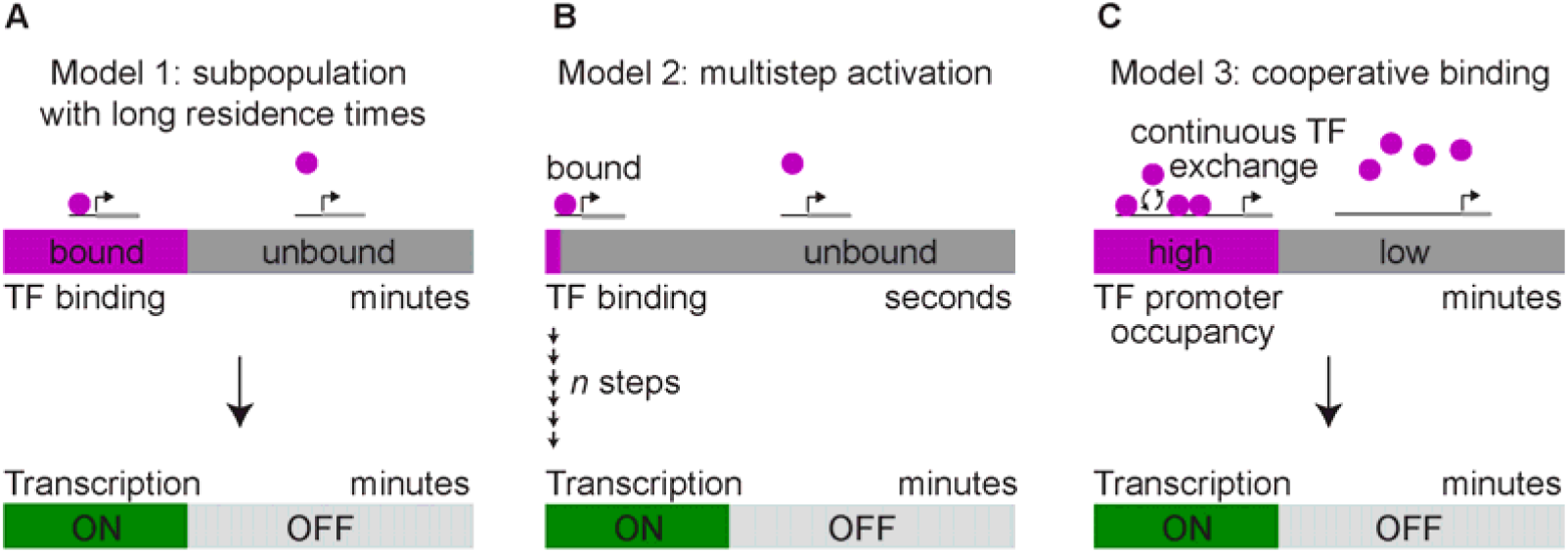
Models to connect short TF binding to long-lasting transcription activation. **A**. A small TF subpopulation may bind with long residence times and activate transcription. **B**. TF binding may initiate a sequence of steps to activate transcription. **C**. Cooperative TF binding may cause longer periods of high and low promoter occupancy, where high occupancy results in transcription.

The most common method to capture TF binding dynamics *in vivo* is single-molecule tracking (SMT). However, interpretation of how detected binding events by SMT activate transcription has been challenging, because these measurements encompass binding to multiple genomic loci with different affinities and are thus a mix of different DNA binding kinetics ^19^. In addition, without imaging transcription of target genes, the functional effect of these binding events is unknown. Although transcription of individual genes can be visualized in living cells by labeling nascent RNA of target gene with the MS2 or PP7 RNA labeling techniques ^20,21^, it has been technically challenging to combine this with SMT. First, TFs have many target genes and with SMT often only a small percentage of molecules is labeled to be able to identify single molecules, thereby significantly decreasing the chance of observing a molecule at the locus of interest. Second, the difference in timescales between TF DNA binding and transcription complicates capturing both timescales without causing photobleaching or phototoxicity. Last, SMT usually only images a single plane inside the nucleus to reduce photobleaching and increase imaging speed, but imaging a single plane limits accurate estimation of the transcription site (TS) intensity and burst duration, as the TS can move out of focus.

To address these challenges, we and others have recently employed 3D orbital tracking to track TSs in 3D and simultaneously measure TF binding and transcription activation of a single target gene in the same cell ^17,18^. These experiments revealed a positive correlation between TF binding and target gene activation, indicating that the fluctuations in TF binding are coupled to the fluctuations in transcription. Moreover, mutations in the TF binding site that reduce the residence time also reduce the burst duration, suggesting a causal link ^18^. However, individual traces measured by 3D orbital tracking were too noisy to extract the functional effect of single TF binding events, preventing in-depth analysis of the kinetic models of transcription activation.

Here, we devised a tracking system to understand how transient TF binding relates to transcription activation. To increase the probability of capturing a sufficient number of binding events, we chose the well-studied yeast TF Gal4 with the highly-expressed target gene *GAL10* as a model system. Gal4 is naturally lowly abundant and only has 15 genomic binding sites and the shared *GAL1-GAL10* promoter is the promoter with the most binding sites (four) ^22^. To measure Gal4 binding and *GAL10* transcription simultaneously, we developed a tracking algorithm to track the *GAL10* TS in a single plane in the *z* direction with an active feedback loop. Using different Gal4 labeling densities and imaging speeds, we tested the different models of transcription activation. We found that the multistep activation model (model 2) was inconsistent with the data, and that binding of the same TF throughout the entire burst (model 1) was infrequent. Instead, our data provide evidence that the main kinetic mode of activation of *GAL10* is that Gal4 molecules show cooperative binding and exchange during a burst (model 3). Our findings suggest a general mechanism for TF-mediated regulation, where TF cooperative binding and TF exchange at multiple binding sites enable prolonged transcriptional bursts.

## Results

### 3D tracking of transcription sites

To measure both Gal4 binding and transcription of its target gene *GAL10* in living yeast cells, we labeled the Gal4 protein with a HaloTag and the *GAL10* nascent RNA with the PP7 system. Shortly, the HaloTag was C-terminally fused to endogenous Gal4 and covalently coupled to JF646 for visualization ^23^. Fourteen PP7 repeats were added to the 5’ end of the *GAL10* gene. Upon transcription, nascent transcripts were bound by PP7 coat proteins fused to GFPEnvy, such that the intensity of the TS reflected the number of nascent *GAL10* transcripts ^21^. Cells were illuminated with a highly-inclined and laminated optical sheet (HILO) to obtain single-molecule sensitivity and to limit photobleaching ^24^.

Measuring both Gal4 binding to the *GAL10* promoter and its effect on *GAL10* transcription requires one to image at high enough acquisition rates to capture TF binding and to image long enough to capture bursting dynamics. In addition, one needs to accurately determine TS intensity. In order to do so, we developed a tracking algorithm to keep the TS in focus with an active feedback loop, while only imaging a single plane. In each frame, the *z* position of the TS was determined using a cylindrical lens ^25^, where the ellipticity of the spot informed on the movement of the TS out of the focal plane. The *z* position of the stage was adjusted accordingly for the next frame. Tracking with only a single plane provides higher acquisition rates and reduces light exposure per time point compared to conventional 3D imaging where several z-planes are imaged, allowing for prolonged total imaging times. Moreover, information of the *z* position of the TS allowed for accurate quantification of the number of nascent *GAL10* transcripts and binding of Gal4 to the locus from a single plane (methods).

We tested how well our refocusing scheme worked by artificially moving a number of fluorescent beads through the focus and recalculating their position from the measured ellipticity (Figure 2A,B). In a range of ^∼^500 nm around the focus (methods), the positions of the beads were reliably determined with a standard deviation of 62 nm (Figure 2C). To limit photobleaching while enabling tracking, we calculated how the acquisition interval affected the total tracking time during which the focus would be kept reliably. Using the measured diffusion speed of the PP7-*GAL10* TS (239 ± 11 nm^2^/s, Figure S1A-B), focus feedback every 5 seconds would theoretically track TSs up to a day (Figure 2D). Without refocusing, the TS would be lost within several minutes (Figure 2D, S1C-D).

**Figure 2.**
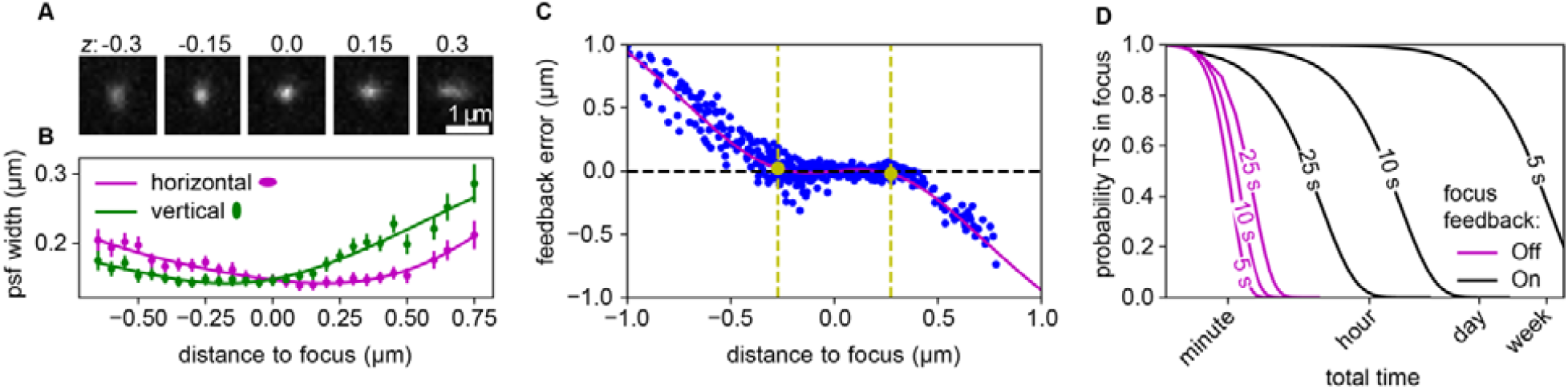
Real-time tracking of TSs using active feedback. **A**. Example images of tetraspec beads imaged at 5 different positions above and below focus at the indicated *z* positions (in μm). The ellipticity of the point spread function (psf) changes with the position. **B**. Width of the psf in horizontal (magenta) and vertical (green) directions at different focus positions. The ellipticity is calculated as the ratio of these. **C**. Error made in focus feedback on a sample with fixed beads as a function of focus position, in a range of 546 nm around the focus (indicated by yellow dotted lines, see methods) the standard deviation of the error was 62 nm. **D**. Probability that the TS is within the depth of focus in all frames in an acquisition. Without focus feedback (magenta), the TS would be lost within minutes, while focus feedback (black) at 25 s, 10 s or 5 s intervals increases the tracking time to tens of minutes, an hour or a day, respectively.

Because the TS intensity fluctuates according to the number of transcripts being produced, the tracking algorithm was adjusted to account for periods of no transcription. If no TS was found, the focal plane was kept constant until the TS re-appeared. Because PP7-*GAL10* is active for >75% of the time ^26^, and the OFF periods are short (exponentially distributed with average ±100 s, Figure S3D,L), the depth of focus of ^∼^250 nm allowed us to track the TS without losing focus with a theoretical probability of ^∼^82%.

### Gal4 binding at *GAL10* reveals a previously undetected long-bound population

We applied this tracking algorithm to measure *GAL10* transcription, while simultaneously monitoring the binding of Gal4 TFs to the *GAL10* locus (Figure 3). Before imaging, yeast was pre-grown in raffinose-containing media, and switched to galactose-containing media to induce transcription of the *GAL10* gene. Upon appearance of a TS, tracking was initiated with feedback every 5 seconds for 20 minutes. In this experiment, Gal4 was labeled densely with JF646 to enable capturing a sufficient number of binding events at our locus of interest. In contrast to SMT, where sparse labeling is required to reliably link molecules between frames ^27^, our method has the advantage of detecting TF binding in the vicinity of a focused target gene without the need for sparse labeling, thereby increasing the TF detection probability significantly. Comparison of the Gal4 intensity distribution of dense versus sparse labeling conditions (Figure S1E) suggested that detected Gal4 binding events with dense labeling represented one or multiple Gal4 molecules, indicating we retain single-molecule sensitivity. Conversely, even in this dense regime, Gal4 labeling efficiency was likely much less than 100% from export of the JF646 dye from yeast cells ^28^. The Gal4-HALO images only showed a few spots in the nucleus (Figure 3A) and did not resemble the images from Gal4-EGFP where all molecules are visible ^29^. We therefor expect that there was additional DNA binding of non-labeled Gal4 molecules at the locus.

**Figure 3.**
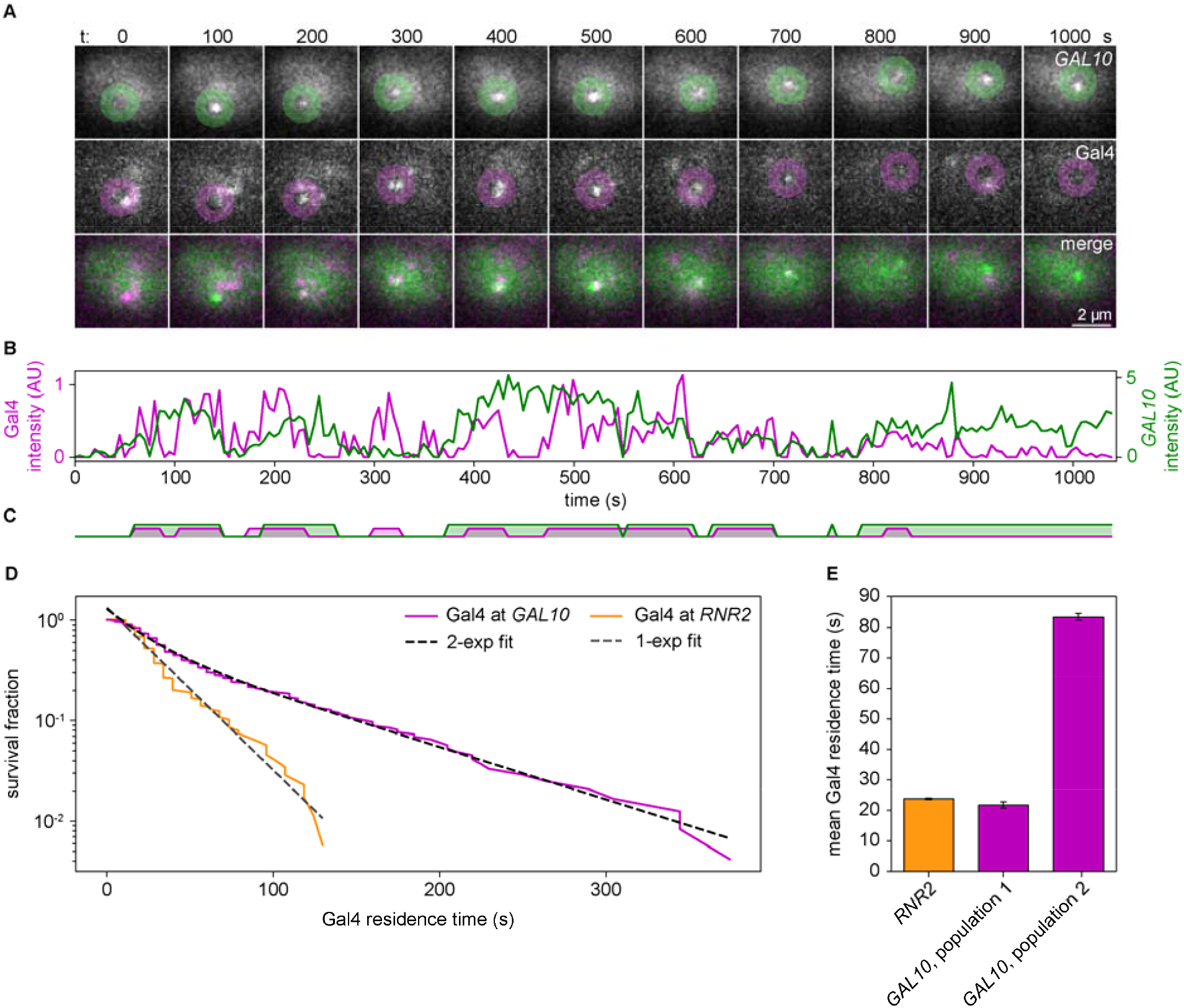
Simultaneous imaging of Gal4 binding and *GAL10* transcription using focus feedback of the *GAL10* transcription site. **A**. Cropped example images of a *GAL10* TS, tracked and imaged every 5 s with focus feedback (top), while simultaneously imaging Gal4 DNA binding at the locus (middle), shown every 100 s. Green and magenta circles indicate the found or interpolated positions of the spot. If no Gal4 spot was found, the position of the *GAL10* TS is shown. Bottom: merge of *GAL10* transcription (green) and Gal4 (magenta). **B**. Example trace of Gal4 binding (magenta) and *GAL10* transcription (green). **C**. Binarization of Gal4 binding and *GAL10* transcription, called by a hidden markov model. **D**. Survival fraction (s/r) of the residence time of HMM-binarized Gal4 residence times at the *GAL10* (magenta) and RNR2 (orange) loci, using dense Gal4 labeling. The probability (s) is corrected for the number of traces (r) for each residence time. Bayes Information Criteria was used to determine whether the survival distribution was best fit with an exponential or a biexponential distribution. Only the best fit is shown in black (for *GAL10*) and grey (for RNR2) dotted lines. For parameters see Figure S3A,E. **E**. Average Gal4 residence times from the exponential fit at RNR2 (orange), and the biexponential fit at *GAL10* (magenta) from the data in D. Error bars represent the standard error of the fit. For D-E: n = 46 cells for Gal4-*GAL10*, n = 34 cells for Gal4-RNR2.

Overall, we obtained traces for 46 cells, where Gal4 binding and *GAL10* transcription was measured simultaneously (Figure 3, Table S1). For periods where *GAL10* transcription was OFF, or where no Gal4 spot was detected within 300 nm, we measured the intensity of the background at the same location. For Gal4, we detected multiple DNA binding events to the *GAL10* gene throughout our 20-min imaging time (Figure 3A-B). The diffraction limit and the physical distance between the PP7-PCP-GFPEnvy label and the TF binding sites made it impossible to resolve whether detected Gal4 molecules in the vicinity of *GAL10* were actually bound to the TF binding site in the GAL1-*GAL10* promoter, or represented background binding, for example to the downstream GAL7 promoter or another nearby gene, or perhaps residing in a Gal4 cluster close to TS ^29^. For simplicity, we refer to all Gal4 molecules within 300 nm of the TS as bound to one of the four Gal4 binding sites in the *GAL10* promoter. We also observed periods without detectable Gal4 binding, which may represent either periods without Gal4 binding or periods during which unlabeled Gal4 molecules were bound.

To characterize the detected Gal4 binding events at *GAL10*, we determined the Gal4 bound periods by binarizing the Gal4 intensity traces with a hidden markov model (HMM) (Figure 3B-C). The distribution of binarized Gal4 binding times at *GAL10* were best fit by a biexponential distribution, indicating two binding populations with different residence times (Figure 3D, S3A). Approximately half of Gal4 binding events (53 ± 1%) lasted on average 22 ± 1 s, and the other half lasted on average 83 ± 1 s (Figure 3E). As a negative control, we also tracked the *RNR2* gene that is not responsive to Gal4. At *RNR2*, Gal4 only showed a single binding population with an average duration of 24 ± 0 s (Figure 3D-E, S3E). Both genes thus displayed binding events of approximately 23 s, which is close to the previously measured 17 s ± 1 s for Gal4 genome-wide binding ^18^. At *GAL10* specifically, we thus captured a previously-undetected second population of binding events with a much longer binding duration, underscoring the potential of our approach to capture longer binding events compared to SMT. The time periods where no Gal4 was detected at both *GAL10* and *RNR2* were exponentially distributed (Figure S3B,F). Although the duration of these Gal4 OFF periods is dependent on the Gal4 labeling density, the observed exponential distributions indicated that the tracking and binarization procedure was not introducing artificial short gaps in Gal4 binding from mistracking or blinking of JF646. Overall, the specific Gal4 binding population also strongly suggested we were able to detect specific Gal4 binding events at the *GAL10* locus.

### Gal4 binding and *GAL10* transcription are dynamically coupled

Using 3D orbital tracking, we previously observed coupling of Gal4 binding and *GAL10* transcription signals with cross-correlation analysis ^18^. To analyze if Gal4 binding and *GAL10* transcription were kinetically coupled in these feedback experiments, we calculated the cross-correlation function between the Gal4 and *GAL10* signals. Similar to our previous results ^18^, the Gal4-*GAL10* cross-correlation showed a peak with a positive delay (Figure S2A). The peak indicates that fluctuations in Gal4 binding and fluctuations in *GAL10* transcription are coupled. The positive delay suggests that Gal4 binding precedes *GAL10* transcription. Transcription at the negative control gene *RNR2* revealed no correlations with Gal4 binding in the cross-correlation (Figure S2B), as expected. These data validate our previous finding that Gal4 binding in the promoter and subsequent transcription of the *GAL10* target gene are temporally coupled.

### Gal4 association and dissociation activates and deactivates transcription rapidly

Having established that we were able to detect specific Gal4 binding at *GAL10* that was coupled to transcription, we aimed to test the different kinetic models of transcription activation (Figure 1). In all three models, TF binding is expected to increase transcription. However, one of the main discriminants between the multi-step activation model (model 2) and the other two models is that in the multi-step model, the TF does not need to be continuously present to activate transcription. Once TFs have initiated the multi-step process to start transcription, they may dissociate while transcription is maintained through the downstream steps. If this multi-step process follows a fixed deterministic time, TF binding may be coupled to the end of transcription, but the delay between TF dissociation and the end of transcription is expected to be long (minutes). Alternatively, if this multistep process is not fixed, but probabilistic – i.e. if transcription is ended by another TF-independent regulatory step – TF dissociation is expected to be completely uncoupled from the end of the transcription event. In both cases, TF dissociation should not be followed by a fast decrease in transcription. In contrast, in the other two models, TF dissociation is expected to decrease transcription.

To discriminate between these predictions, we aligned the Gal4-*GAL10* time traces to the starts (Gal4 association) and ends (Gal4 dissociation) of all detected HHM-detected Gal4 binding events and plotted the average intensity of the Gal4 signal and of the *GAL10* TS (Figure 4A-B). As a consequence of the alignment process, aligning to the start of Gal4 binding displayed a Gal4 signal increase from timepoint 0. The number of mRNAs at the TS increased without detectable delay (5 s time resolution), indicating that Gal4 association has an almost immediate effect on transcription. Alignment to the end of the Gal4 binding events revealed that TF dissociation also coincided with a decrease in transcription (Figure 4B). The negative control gene *RNR2* did not show any transcriptional response with this analysis (Figure S4A-B). These findings suggest that Gal4 binding determines both the start and the end of a transcriptional burst, which would be in line with models 1 and 3, but not with the second multi-step model, where TF binding only determines the start, but not the end of a burst.

**Figure 4.**
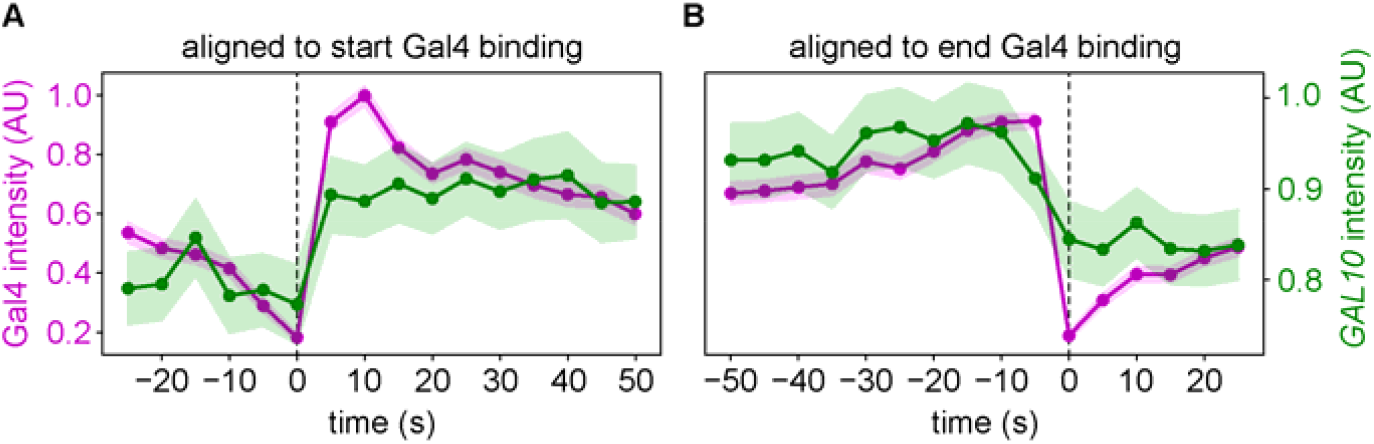
TF binding and release determine burst start and end. **A**. Average Gal4 (magenta) and *GAL10* TS (green) intensities aligned to the start of Gal4 binding. **B**. Same as A aligned to the end of Gal4 binding. Shaded areas indicate standard errors of the means.

Because the effect of Gal4 DNA binding on transcription was already detected within 5 seconds, the experiment was repeated with a faster time interval of 1 second. With this higher time resolution, we observed that transcription increased already after 1 second of Gal4 association, and was maximally increased after 2 seconds (Figure S4C). This short delay may be an underestimation of the true delay because we may have been imaging in the wrong focal plane before the TS was detected and therefore may have missed prior Gal4 binding. However, release of Gal4 from an accurately traced TS also causes a decrease in transcription within a second (Figure S4D), which argues against a large contribution of such missed events. This short delay between Gal4 dissociation and transcription decrease may be surprising given the elongation time of Pol II molecules along the gene, but can be possible if the *GAL10* gene is completely saturated with Pol II molecules. In that case, loss of transcription initiation immediately causes a reduction in the PP7 signal from the release of transcripts from Pol II molecules that already reached the 3’ end. This complete Pol II coverage occurs when the burst duration is longer than the residence time of a single Pol II during elongation and termination, which is indeed in line with the measurements at *GAL10* (burst duration: 138 ± 1 s (Figure S3C), estimated residence time of a single *GAL10* transcript: ^∼^47 s ^26^). Overall, our data suggests that Gal4 association and dissociation causes an increase and decrease in transcription within a few seconds, respectively.

Lastly, to further validate this rapid activation, we selected the 25% shortest binding events, binding for less than 4 seconds, and found that these were indeed able to weakly increase transcription (Figure S4E-F). We therefore conclude that Gal4 binding acts on transcription rapidly and thereby determine the start and end of transcriptional bursts, which is inconsistent with the multistep activation model.

### Transcription activation by Gal4 is at least 20% efficient

The above analysis indicates that Gal4 binding is followed by transcription activation on average, but not every binding event is necessarily productive. To understand how efficient transcription activation is, we determined how many binding events resulted in a more-than-random increase in the *GAL10* intensity. We calculated the derivatives of the *GAL10* transcription trace, revealing whether transcription increases or decreases at all timepoints. The traces were then aligned at the start of Gal4 binding events, which showed that TF binding resulted in a more-than-random increase in transcription in 21 ± 6% of the traces (Figure 5A, methods). Because likely many detected Gal4 binding events may not have actually bound to *GAL10*, but may have represented background binding to other loci in the vicinity of the *GAL10* gene, this ^∼^20% efficiency represents a lower bound (see discussion).

**Figure 5.**
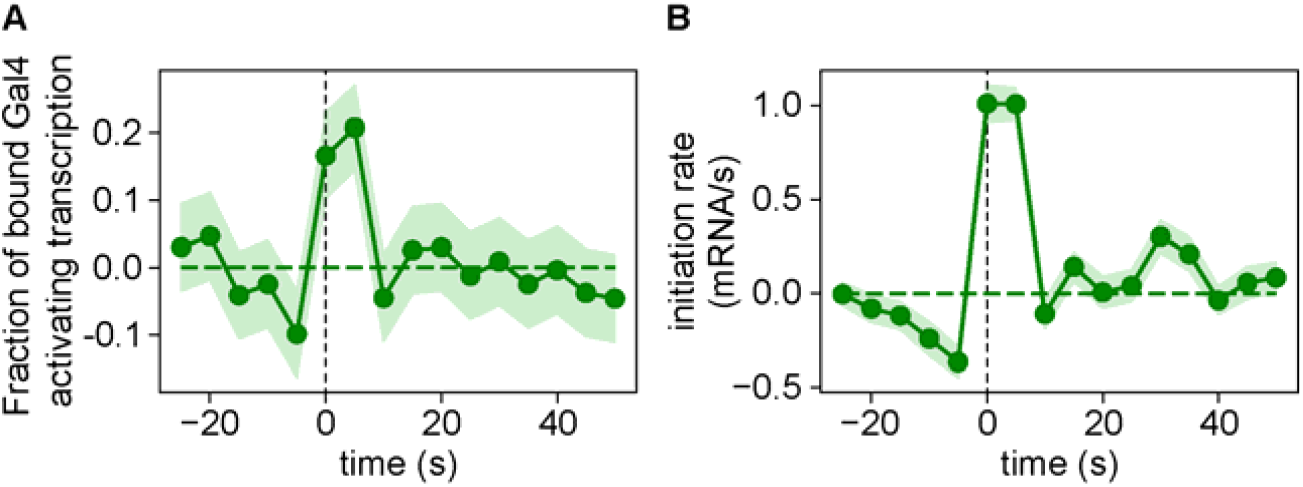
The TF Gal4 activates transcription efficiently. **A**. Fraction of Gal4 binding events resulting in transcription activation, as measured by the faction of *GAL10* TS intensities increasing in intensity, corrected for random chance. Data is aligned at the start of Gal4 binding events. **B**. *GAL10* initiation rate, as measured by the average increase of TS intensity. The measured *GAL10* intensity was converted into number of nascent *GAL10* transcripts as described in the methods. Data is aligned at the starts of *GAL10* bursts. Shaded areas indicate standard error of the mean.

This 20% transcription activation efficiency indicates how often a transcriptional burst starts after a TF binds to DNA. To determine the efficiency of pol II loading once a burst has started, we calculated the rate of transcription initiation. First, the measured *GAL10* intensity was converted into number of nascent *GAL10* transcripts by normalizing the distribution of *GAL10* intensities to the PP7-*GAL10* nascent mRNA distribution, obtained from smFISH ^26^ (methods, Figure S5A). Second, the *GAL10* intensity traces were binarized with a HMM, and were aligned at the start of each transcriptional burst (Figure 5B). The slope of the transcription increase represented the average initiation rate, which peaked at 1.0 ± 0.1 mRNA per second. The increase from zero to the maximum initiation rate was reached within the first 5 seconds, suggesting that polymerases reached maximum efficiency quickly. To ensure this rate was not affected by possible tracking errors at the beginning and ends of bursts, we repeated this analysis by tracking a DNA label (tetO/tetR-tdTomato) that was inserted near the *GAL10* gene ^30^, and measuring *GAL10* transcription in a second channel, and we found a very similar rate of 1.1 ± 0.1 mRNA/s (Figure S5B). The initiation rate of ^∼^1 mRNA/s is very similar to the delay of the first transcription initiation event after Gal4 binds to the locus, suggesting that the loading of the first and subsequent RNA polymerases are on comparable timescales. Once Gal4 binds, RNA polymerases are thus rapidly loaded onto the gene. Overall, these analyses indicate that TF association starts a burst at least 20% of the time and then load up to ^∼^1 polymerases every second.

### Gal4 exchanges during transcriptional bursts

The findings that TF association and dissociation increases and decreases transcription, respectively, suggest a model where TF occupancy is required throughout the burst, which is consistent with both model 1 and 3. TFs may bind throughout the entire burst, such that the TF residence time directly determines the burst duration (model 1). In addition, since the *GAL10* promoter contains four Gal4 binding sites, individual molecules may also exchange during a burst (model 3). These two models are not mutually exclusive, since some molecules could remain bound during the entire burst, whereas other molecules could exchange.

To investigate the contribution of these two kinetics modes of activation at *GAL10*, we analyzed whether there was evidence for TFs exchange during a burst (model 3). For both models (1 and 3), TFs are expected to associate at the beginning of the burst and dissociate at the end. In addition, in the cooperative binding model, TF molecules are expected have a high frequency of association and dissociation during a burst as well, as individual TF molecules exchange. Since not all Gal4 molecules are labeled, these exchange events between labeled and unlabeled Gal4 should be detectable in the traces. To test these predictions, we determined ON and OFF periods of *GAL10* with a HMM, and classified five periods: before a burst, at the start of a burst, during a burst, at the end of a burst, and after a burst (Figure 6A). The occupancy of Gal4 was higher during a *GAL10* burst than during all other periods (Figure 6B), providing confidence that the burst periods were determined correctly and suggesting that continuous Gal4 presence is required for transcription activation. Gal4 occupancy was not significantly different or even lower during a burst than during other periods for the negative control gene *RNR2* (Figure S6A).

**Figure 6.**
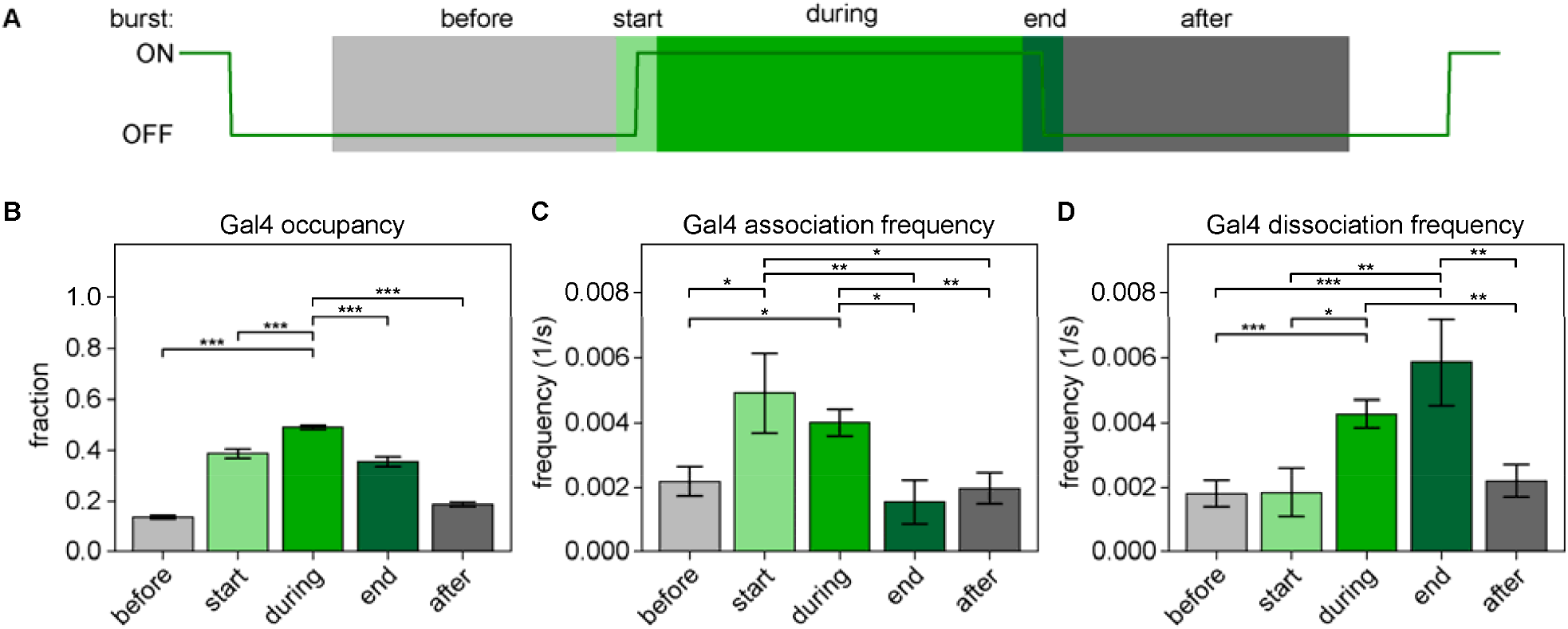
Gal4 molecules exchange during a transcriptional burst. **A**. Classification of the five periods: before (>25 s after previous burst – 10 s before start burst), at the start (10 s before – 10 s after start burst), during (10 s after start – 10 s before end burst), at the end (10 s before – 10 s after end burst), and after (10 s after burst – >25 s before next burst) a burst (green line). **B**. Gal4 occupancy at *GAL10*, as measured by the fraction of timepoints with bound Gal4, in the five periods indicated in A. **C**. Gal4 association frequency at *GAL10*, as measured by the number of Gal4 association events divided by number of frames, in the five periods indicated in A. **D**. Gal4 dissociation frequency at *GAL10*, as measured by the number of Gal4 dissociation events divided by the number of frames, in the five periods indicated in A. In C and D only binding events lasting longer than 20 seconds are considered. Error bars in B-D indicate standard error of the mean. In B, significance is only calculated for “during at burst” versus all other periods, in C and D, significance was calculated between all time periods, and only significant bars are shown. Significance was determined by bootstrapping with 100.000 repeats, and corrected for multiple testing using the Bonferroni method. ^*^ *p* < 0.05, ^**^ *p* < 0.005, ^***^ *p* < 0.0005

When determining the association and dissociation frequency of Gal4 in these five periods, we observed a weak tendency for Gal4 molecules to associate with a higher frequency at the start of a burst (Figure S6B) and to dissociate with a higher frequency at the end of the burst (Figure S6C), as expected, but these differences were only weakly or not significant. We therefor only selected binding events longer than 20s, reasoning that this may enrich for *GAL10*-specific binding events, since *GAL10* displayed a specific population of long-bound molecules (average residence 83 ± 1 s, Figure 3D-E, S3A) that was not present at *RNR2* (Figure 3D-E, S3E). Upon selecting for these long-bound molecules, Gal4 association and dissociation were clearly enriched at the starts and ends of bursts, respectively, compared to periods before or after a burst (Figure 6C-D). Moreover, as predicted by the TF exchange model, the Gal4 association frequency was significantly higher during a burst than before or after a burst, and equally high as at the start of a burst. Similarly, Gal4 dissociation frequency was as high during a burst as at the end of a burst. Although apparent Gal4 dissociation could technically also arise from photobleaching rather than Gal4 dissociation, the equally high frequency of Gal4 association argues against such technical confounding effects. Gal4 displayed similar increased association and dissociation rates during *GAL10* bursts compared to other periods when tracking with 1 s interval (Figure S6D-E), but not at the negative control gene *RNR2* (Figure S6F-G). Overall, these high Gal4 association and dissociation frequencies during a burst are in line with Gal4 molecules exchanging during a burst and thus support the cooperative binding model (model 3).

### Gal4 rarely binds throughout the entire burst

Next, we investigated the contribution of model 1, where Gal4 molecules remain bound during the entire burst. Comparison of the distribution of the Gal4 bound periods to the distribution of the *GAL10* ON periods revealed that the average *GAL10* burst duration (138 ± 1 s, Figure S3C) was almost twice as long as the long-bound TF population (83 ± 1 s, Figure S3A), suggesting that the majority of molecules bind with shorter residence times than the burst duration. However, the higher labeling density in these measurements implied that multiple Gal4 molecules could be bound at the same time. To be able to determine the residence time of single Gal4 molecules, we repeated the Gal4-*GAL10* tracking experiment with sparsely labeled Gal4. A 5 s time interval was used to limit photobleaching effects. We note that, at this interval, it may be possible that two Gal4 molecules measured in consecutive frames may in fact have exchanged in between the frames and therefore do not belong to the same binding event. However, since on average less than one Gal4 molecule was labeled per cell, we assumed that consecutively measured molecules in the same location belonged to the same binding event. As expected, the Gal4 intensities from sparse labeling followed a narrow unimodal distribution (Figure S1E), confirming we detected single rather than multiple Gal4 molecules. In these experiments, a DNA label was also inserted near the *GAL10* gene to confirm we reliably tracked during transcriptionally inactive periods ^30^.

Similar to the experiments performed with dense labeling, the resulting survival probability distribution of the Gal4 residence times at the *GAL10* locus was best described by a biexponential distribution, indicating two binding populations, where 82% of Gal4 bound on average for 14 ± 0 s and 18% of Gal4 bound on average for 89 ± 1 s (Figure 7A, S3I). These residence times were close to the residence times found with the dense labeling (Figure 3D-E, S3A), suggesting that binding multiple Gal4 molecules at *GAL10* follows similar kinetics as binding of single molecules. In the sparse labeling experiment, the *GAL10* burst duration was slightly longer (169 ± 1 s, Figure S3K) than measured with the dense labeling (138 ± 1 s, Figure S3C), perhaps because tracking of the DNA label rather than the TS directly resulted in better tracking in the beginning or end of bursts.

**Figure 7.**
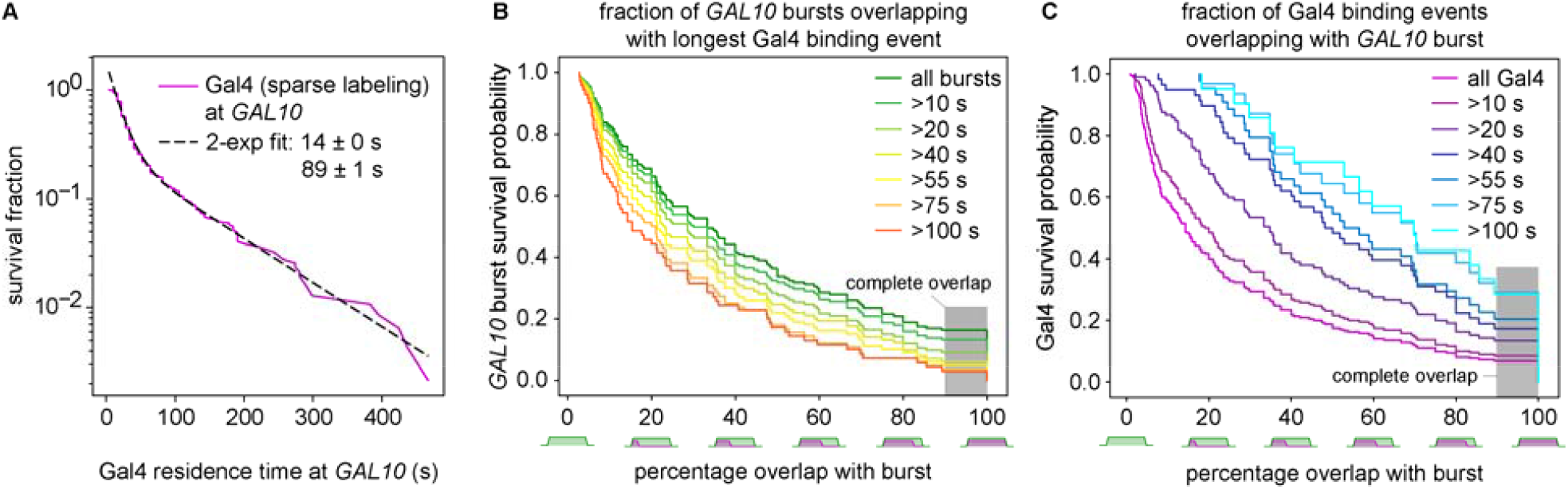
Gal4 rarely binding throughout the entire burst. **A**. Survival fraction (s/r) of the residence time of individual Gal4 molecules at the *GAL10* locus, determined using sparse Gal4 labeling. The probability (s) is corrected for the number of traces (r) for each residence time. Black dotted line represents the biexponential fit to the measured Gal4 residence times (magenta) with the indicated parameters. Errors indicate standard error of the fit **B**. Survival probability of the fraction of *GAL10* bursts (y-axis) that overlap with the longest Gal4 binding event for at least the indicated percentage of the burst (x-axis), plotted for bursts with different lengths (colored lines). **C**. Survival probability of the fraction of Gal4 binding events (y-axis) overlapping with a *GAL10* burst with the indicated percentage (x-axis), plotted for Gal4 binding events of different lengths (colored lines). Grey shading indicates the threshold of 90% overlap to define complete overlap of a Gal4 binding event with a burst.

To understand whether Gal4 molecules stay bound throughout the entire burst, we calculated the percentage of the burst that overlapped with a Gal4 binding event (Figure 7B). Bursts that did not overlap with any Gal4 binding event were not included in this analysis. Since the HMM binarization may introduce errors in defining the starts and ends of binding events and bursts, complete overlap of a burst with a Gal4 binding event was defined as bursts that overlapped with Gal4 for 90% of the burst or longer (grey areas, Figure 7B-C). We find that approximately 16% of the bursts overlapped completely with a Gal4 binding event (Figure 7B). Of the long bursts (>100s), only 3% showed complete overlap with Gal4 binding, suggesting that these long bursts arise from TF exchange. Moreover, the same analysis from the perspective of the Gal4 TF revealed that only 7% of all Gal4 binding events overlapped completely with a *GAL10* burst (Figure 7C). When selecting for longer Gal4 binding events (>100s), the percentage of binding events with complete overlap increased to 29%. Longer binding events are thus more likely to completely overlap with a burst. However, these results also indicate that although Gal4 can bind for the entire burst, that this occurs only infrequently. We therefore conclude that the main kinetic mechanism of Gal4 binding is exchange of Gal4 molecules during a burst.

### TF exchange increases the burst duration

To further validate that the cooperative binding model is the main kinetic mode of activation, we calculated the number of binding events per burst. If there is no exchange (model 1), then every burst is expected to overlap with one binding event, irrespective of the duration of the burst. If, however, TFs exchange during a burst (model 3), then longer bursts are expected to show more Gal4 binding events. Since any stochastic process, even background binding, would result in more binding events for longer bursts, the number of Gal4 binding events at *GAL10* were compared to the number of binding events at *RNR2* as a control. Without TF exchange, *GAL10* and *RNR2* are expected to show similar numbers of binding events, whereas Gal4 exchange should show an increase at *GAL10* compared to *RNR2* for longer bursts. The absolute number of binding events depends on the labeling percentage, but the relative number can be compared between bursts of different durations. We divided the bursts in two equal halves representing short and long burst durations and plotted the number of TF binding events per bursts (Figure S7). We observed significantly more Gal4 binding events for longer bursts at *GAL10* than at *RNR2*, of which the latter even had a higher Gal4 labeling density, supporting the model that TFs exchange during a burst. Conversely, the findings that long burst show multiple binding events (Figure S7) and only rarely overlap completely with a single TF binding event (Figure 7B), also suggests that TF exchange prolongs the total period of TF occupancy and the total burst duration.

## Discussion

In this study, we developed a tracking algorithm to simultaneously measure Gal4 binding at the *GAL10* gene and the resulting effect on *GAL10* transcription. This method allowed us to test the different kinetic models for TF-driven transcription activation. We find that Gal4 occupancy is required during the entire burst, where association and dissociation of Gal4 rapidly results in activation and inactivation of transcription, respectively. Although some Gal4 molecules stay bound throughout the entire burst, the majority of Gal4 molecules exchanges during a burst, which increases the duration of the TF bound periods and the burst duration. Our results thus support the model where the timescales of short TF binding dynamics and longer transcriptional bursts are connected through cooperative binding of TFs.

### TF exchange at a gene with multiple binding sites

In a previous study, we found that lowering the affinity of the single binding site in the GAL3 promoter reduced the burst duration^18^. This reduced burst duration correlated with reduced Gal4 DNA residence time *in vitro*, strongly suggesting that Gal4 residence time directly determines the burst duration at genes with a single binding site. At the *GAL10* gene with multiple Gal4 binding sites, the current study suggests that residence time is only directly linked to the burst duration in maximally 16% of all bursts, but that the majority of bursts shows Gal4 exchange. Rather than describing promoters as TF bound or TF unbound, promoters with multiple binding sites may be better described by high and low/no TF occupancy states. During the high occupancy state, individual TFs exchange at the different binding sites within the same promoter, while maintaining high promoter occupancy. TF exchange may increase the total time of the high occupancy state, and consequently the burst duration. We expect that the time of high TF occupancy is correlated with the TF residence time, since longer TF residence times likely decrease the probability of ending the high occupancy state. This possible coupling between TF residence time and high occupancy periods may explain why previous reports found that transcriptional output is predicted well by TF residence times ^31,8,17,16,10^, even though residence times and transcriptional bursts are on different timescales.

### TF cooperativity as a potential general kinetic activation mode

The finding that Gal4 molecules show higher DNA association rates during a burst than in between bursts suggests that instead of independent binding, Gal4 molecules bind to multiple target sequences cooperatively ^12^. In support, previous studies have reported cooperative binding of Gal4 in vitro and *in vivo* ^32,33^. Recently, we showed that Gal4 forms clusters ^29^, and that self-interactions within a cluster enable recruitment of additional Gal4 molecules to sites were Gal4 is already bound to target genes, which may provide a mechanism for cooperative binding. Additionally, TF cooperativity and increased TF binding during a burst could arise from nucleosome displacement ^34^, stabilization of TFs on the DNA ^15^, or DNA shape ^35^. Moreover, mechanisms that inhibit TF binding during inactive periods can also contribute to the apparent cooperativity, for example by creating promoter chromatin states that are non-permissive for TF binding ^12,36^.

Since many TFs show cooperative binding ^37^ and many promoter and enhancers contain multiple TF binding sites, cooperative binding and exchange of TFs may turn out to be a general kinetic mode to bridge the timescale gap between short dwell times of DNA-bound factors and longer transcription activation. This cooperative model is not restricted to single TFs, but may include cooperation by multiple different TFs. Single-molecule footprinting studies in *Drosophila* and mouse cells found evidence for cooperative TF binding at regulatory regions *in vivo*, which was largely independent of TF identity ^38,39^. The tracking method developed in this study will be an essential tool to understand the kinetic principles how multiple TFs cooperate at promoters and enhancers to regulate gene expression in more complex eukaryotes. Since TSs move slower in human cell lines (71 ± 5 nm^2^/s in U2Os cells, Figure S1A) the interval between the frames may even be prolonged compared to yeast (to 20-30 seconds) without losing focus. However, the current setup may be suboptimal to detect sufficient numbers of TF binding events for genes that are less transcriptionally active, as is often the case for mammalian genes, but we expect that future improvements will enable wider applications to test the generality of our findings across organisms.

### Transcription efficiency

We find that Gal4 binding at *GAL10* starts a burst with 21 ± 6% efficiency. This number is likely a lower bound, since our classification of Gal4 binding events also included Gal4 molecules that were in fact bound to different loci that were in the proximity of the *GAL10* gene. To estimate the “true” efficiency, we attempted to correct for these detected non-*GAL10* binding background binding events. We speculated that the frequency of Gal4 binding to the *RNR2* locus was a good approximation for the background binding frequency at *GAL10*. Comparison of the Gal4 association frequency at the start of the *RNR2* and *GAL10* bursts (3.5·10^−3^ ± 1.1·10^−3^ s^-1^ for *RNR2* versus 6.1·10^−3^ ± 1.3·10^−3^ s^-1^ for *GAL10*) suggested that approximately 57 ± 21% of Gal4 binding events at *GAL10* represented background binding. When adjusting for this background binding, we estimate that Gal4 activates *GAL10* transcription with 49 ± 29% efficiency. Gal4 thus appears to efficiently activate transcription.

In addition, we measured maximum initiation rate of ^∼^1 mRNA/s at the beginning of the burst. This rate appears higher than the previously estimated maximum initiation rate of 0.13 — 0.17 mRNA/s at HIS3 in yeast ^40^, although we note that we only detect this high rate for the first ^∼^10 s of a burst (Figure 5B, S5). Considering that the initiation rates may be lower than 1 mRNA/s during the rest of the ON periods, and that *GAL10* has OFF periods without initiation, the effective average initiation rate is likely in the same range as HIS3. However, other studies have suggested initiation rates of two orders of magnitude lower: 0.001 – 0.02 mRNA/s or 0.01 PIC/s for the average yeast gene and 0.006 mRNA/s for *GAL10*^41,42^. Such low initiation rates would result in an occupancy of less than 1 nascent transcript at the *GAL10* transcription site, which is inconsistent with both the live-cell and smFISH measurements at *GAL10*. Recently, single-molecule imaging of PIC components *in vivo* suggested that PIC assembly at the average yeast gene takes ^∼^5 s ^42^. Given that *GAL10* is much higher transcribed than the majority of the yeast genes ^43^, our estimation of 1 s per PIC assembly at *GAL10* would be conceivable.

Both the efficiency of starting a burst and the initiation rate during a burst suggest that transcription efficiency of the *GAL10* gene in yeast is very high. In mammalian cells, previous *in vivo* FRAP-based measurements have suggested that only 1% of Pol II molecules results in a full-length mRNA ^44,45^. In our experiments, the efficiency of Pol II itself is unclear and, in principle, it is possible Pol II has many failed attempts before entering productive elongation, observable by an increase in the PP7 signal. With the measured *GAL10* initiation rate of ^∼^1 mRNA/s, a 1% Pol II efficiency would imply a rate of Pol II loading of 100 Pol II molecules/s, which may not be very likely. We therefore favor the explanation that transcription in budding yeast is more efficient than in more complex eukaryotes. In line with this, the measured maximum initiation rate of ^∼^1 mRNA/s at *GAL10* appears to be higher than the estimated rates for *Drosophila* (0.07 – 0.17 mRNA/s)^46,47^, and for mammalian cells (0.008 – 0.03 mRNA/s)^44,48^. Besides reduced rates of promoter escape, more complex eukaryotes have additional regulation at pause release, which is not present in budding yeast, and which may reduce the overall transcription efficiency ^45^.

### Gal4 binding populations

The distribution *GAL10* residence times revealed two binding populations with residence times of ^∼^ 15-20 s and ^∼^ 85-90 s. The latter binding population was not detected previously, even when attempting to measure residence times in the vicinity of the *GAL10* transcription sites^18^, suggesting that residence times by conventional SMT are restricted by for example photobleaching and by movement of genes out of the focal plane. The shorter binding population of 15-20 s is much longer than the residence time typically found for non-specific binding (^∼^ 1 s)^28,18^. Such 1 s non-specific event would not be captured with our 5 s time-interval. Since a similar 15-20 s residence time was also found at the negative control gene *RNR2*, and for Gal4 genome-wide binding ^18^, we argue that these binding events are a combination of specific binding at *GAL10* and specific binding at other nearby genes. The idea that these shorter binding events are specific is supported by our result that Gal4 binding events lasting less than 4 s could still weakly activate transcription (Figure S4E-F). Perhaps these shorter binding events represent binding of single Gal4 molecules to a single site, without cooperative binding at multiple sites. Future studies with promoter mutations in the Gal4 binding sites will provide insight into these binding populations further.

In summary, by simultaneously measuring transient TF binding to a single locus and the effect of these binding events on transcription, we uncovered that TF cooperativity and TF exchange connect transient TF binding to prolonged transcription activation. We speculate that enhancers, which contain multiple TF binding sites, employ the same cooperative mechanisms to create prolonged high TF occupancy states that promote initiation of large transcriptional bursts. Given the widespread occurrence of TF cooperativity from single-cell to more complex organisms, our results suggest that TF exchange may provide a widely-used mechanism to robustly activate gene transcription.

## Acknowledgements

We thank D. Mazza and E. de Wit for critical reading of the manuscript and F. van Leeuwen, A. Coulon and members of the T.L.L lab for helpful feedback and suggestions. We thank Daniel Larson for U2OS cells, Linda Joosen for assistance with the growth of U2OS cells, Luke Lavis for the dyes JF646 and JFX650, and Amir Aharoni for plasmids. We thank the Research High Performance Computing Facility of the NKI for assistance. This work was supported by an institutional grant of the Dutch Cancer Society and of the Dutch Ministry of Health, Welfare and Sport, the Dutch Research Council (gravitation program CancerGenomiCs.nl), Oncode Institute, which is partly financed by the Dutch Cancer Society, and the European Research Council (ERC Starting Grant 755695 BURSTREG).

## Materials and Methods

### Yeast strains and plasmids

All BY4743 diploid yeast strains (*Saccharomyces cerevisiae*) used in this study were derived from mating a BY4741 and a BY4742 haploid parent strain, in which endogenous *GAL4* was fused to *HALO* and *PDR5* was deleted. To create *pdr5Δ::HPH*, strains were transformed with a PCR product from plasmid pTL083 containing the Hygromycin B (HPH) resistance cassette which integrated at *PDR5*. To create *GAL4-HALO*, strains were transformed with a PCR product from plasmid pTL088 containing the *HALO-loxP-kanMX-loxP* cassette which integrated at the 3’-end of *GAL4*, followed by *kanMX* removal with CRE recombinase from pTL014.

The PP7 loops required to visualize transcription were integrated in one of the haploid parent strains prior to mating. To integrate fourteen PP7 loops, a PCR product from plasmid pTL031 containing the PP7 loop cassette and *loxP-kanMX-loxP* was integrated at the 5’-end of either *GAL10* or *RNR2*, followed by *kanMX* removal with CRE recombinase from plasmid pTL014. PP7 coat proteins (PCP) were either expressed from plasmid pTL092 or integrated in one of the parent strains using PacI-digested plasmid pTL174.

To enhance the HaloTag labeling efficiency, besides *PDR5*, another exporter was deleted: *YOR1*. To create *yor1Δ* in both parent strains, a CRISPR-based approach was used [5]: strains were co-transformed with a URA-plasmid expressing Cas9 and a guide RNA targeting *YOR1* (pTL407), together with a single stranded oligo (5’-TATTACTGTTTTTATATTCAAAAAGAGTAAAGCCGTTGCTATATACGAATTTTATATTATTTGTTGCATGATTTTTCTCTTTTATTTATTTATATGTTGC-3’) as repair template, whereafter the Cas9 plasmid was removed using 5-Fluoroorotic Acid (5-FOA) selection.

The DNA label was introduced at the *GAL* locus using three successive rounds of transformations, as described in ^30^. Firstly, a PCR product encoding natMX flanked by homology arms for the tetO array was integrated at 3’-*GAL1*. Secondly, a similar CRISPR approach was used as described above, with a Cas9 plasmid targeting the natMX cassette (pTL552) and using a PCR product containing the *tetOx128* array as double-stranded repair template. Finally, *tetR1-tdTomato* was integrated at the *ADE1* locus by transformation using a plasmid digestion as a repair template. All integrations were checked with PCR. Cells were grown at 30°C in synthetic media. Strains and plasmids used to construct the strains can be found in **Tables S1 and S2**, respectively.

**Supplementary Table 1:** Yeast strains used in this study.

| Strain | Genotype | Source |
| --- | --- | --- |
| YTL431 | BY4743 with *pdr5::HPH/pdr5::HPH*, *GAL4-HALO/GAL4-HALO* (w/o *kanMX*), *RNR2/14xPP7-5'-RNR2* (w/o *kanMX*) + pTL092 | <sup>18</sup> |
| YTL639 | BY4743 with *pdr5::HPH/pdr5::HPH*, *GAL4-HALO/GAL4-HALO* (w/o *kanMX*), *GAL10/14xPP7-5'-GAL10* (w/o *kanMX*) + *ura3Δ0/ura3Δ0::pRPL15A-PCP-GFPEnvy* | This study |
| YTL1649 | BY4743 with *pdr5::HPH/pdr5::HPH*, *GAL4-HALO/GAL4-HALO* (w/o *kanMX*), *GAL10/14xPP7-5'-GAL10* (w/o *kanMX*) + *ura3Δ0/ura3Δ0::pRPL15A-PCP-GFPEnvy*, *yor1Δ/yor1Δ*, *GAL1/tetOx128-3'-GAL1*, *ADE1/ade1::tetR1-tdTomato-kanMX* | This study |

**Supplementary Table 2:**
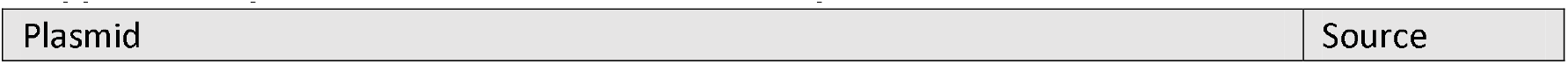
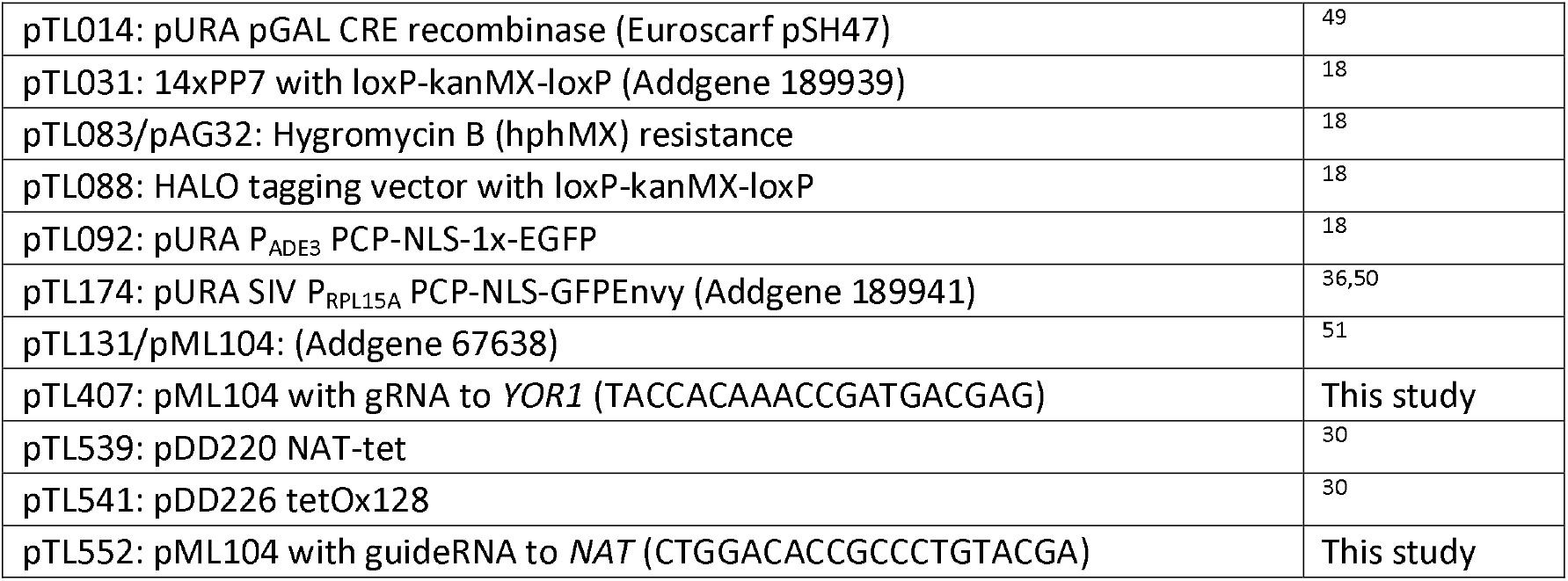
Plasmids used in this study.

### Microscopy

Yeast cultures were grown to early mid-log in synthetic complete (SC) medium with 2% raffinose, washed with medium containing 2% galactose and imaged on a coverslip with a 2% agarose pad containing 1% raffinose and 2% galactose for induction of *GAL10*, as described previously ^52^. For YTL431, cells were grown in SC medium without uracil and 0.005% methyl methane sulfonate for induction of *RNR2*. Cells were incubated with 20 nM Janelia Fluor 646 ^23^ for two hours (YTL639, YTL431) or 0.5 nM Janelia Fluor X 650 ^53^ for 15 minutes (YTL1649) before imaging in order to label Gal4 densely or sparsely, respectively. All samples were mounted on washed and sonicated Zeiss Coverslip HI (000000-1787-996) coverslips, as described previously ^52^.

Imaging was performed on a Zeiss AxioObserver.7 / ELYRA.P1 microscope equipped with an incubator for microscopy (Pecon Incubator XL Tirf Dark S1 Nano) set at 30°C, Zeiss Scanning Stage Piezo 130×100 and Coherent Sapphire 405, 488, 561 and 640 nm lasers with maximum powers at 50, 100, 100 and 150 mW respectively. Images were acquired with a Zeiss alpha Plan-Apochromat 100× numerical aperture (NA) 1.57 oil objective with Immersol HI 661 immersion oil and a 1.6× Zeiss optovar, yielding a 97.09 nm pixel size. For two-channel experiments (YTL639 and YTL431) a Zeiss LBF 405/488/642 filter set was used and the emission was split in two channels using a Zeiss duolink splitter holding a filter set with a Zeiss BS561 dichroic beamsplitter and Semrock FF03-525/50-25 and BLP02-561R-25 emission filters and imaged simultaneously on two Andor EM-CCD iXon DU 897 camera’s. For three-channel experiments (YTL1649), a filter set consisting of a Chroma ZT405/488/561rpcv2-UF1 dichroic filter and a ZET405/488/561/640mv2 emission filter was used, and the emission was imaged subsequently without delay between the channels on one of the cameras.

All imaging was performed using Zen Black 2.3 and the following settings: TIRF_uHP acquisition mode, 256×256 pixels field of view, 1.6x optovar, highly-inclined and laminated optical sheet (HILO) illumination mode, 100 ms exposure time, EMCCD gain set to 100×, imaging interval: 1 s or 5 s. Gal4-JF646 was imaged with 640 nm excitation at 0.4% (of ±20mW) power resulting in ±100 W/cm^2^ excitation intensity (YTL639, YTL431) or at 1% power resulting in ±250 W/cm^2^ (YTL1649). The *GAL10* or *RNR2* TS was imaged with 488 nm excitation at 0.01% (of ±25 mW) power resulting in ±3 W/cm^2^ excitation intensity. The DNA label-tdTomato (YTL1649 only) was imaged with 561 nm at 0.1% (of ±25 mW) power resulting in 30 W/cm^2^ excitation intensity.

### MSD calculations

To calculate the mean square displacement (MSD) of transcription sites in yeast cells and human cell lines with respect to the microscope we acquired several z-stacks in timelapse. Yeast cells (YTL639) were cultured and induced as described before and human cells (U2OS with a MS2 containing reporter gene driven by a 85x PonA responsive promoter ^54^) were cultured as described by ^54^ and stimulated with Ponasterone A for 24h prior to imaging.

The *z-*stacks had an interval between 250 and 600 nm and 9 to 20 slices and in timelapses with up to 300 frames between 2.5 and 6.5 seconds apart. For the U2OS cells, a Zeiss alpha Plan-Apochromat 100x/1.46 Oil objective was used with 0.04% 488 nm laser (±12 W/cm2) and 20 ms exposure. For the yeast cells, a Zeiss alpha Plan-Apochromat 100x/1.57 Oil-HI objective was used with 0.02% 488 nm laser (±6 W/cm^2^) and 30 ms exposure.

The images were segmented into cell and nucleus 3D label masks with a Python script using Otsu thresholding and watershedding. Candidate particles were found in the 3D images using a Laplacian of Gaussian filter with sigma 1.67 pixels. These candidate particles were then fit with

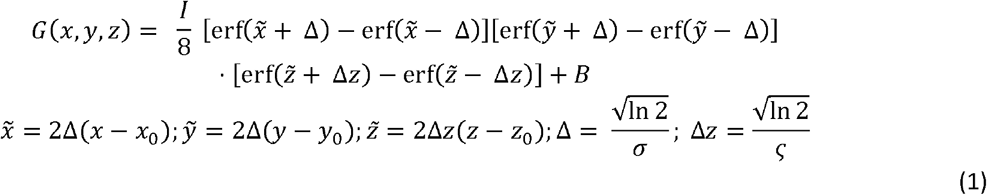

Here (*x*_0_, *y*_0_, *z*_0_) is the center position of the particle, *I* its intensity, *B* the background intensity and σ and ς the widths of the peak in *xy* and *z* respectively. Subsequent filtering on localization precision, peak width and R^2^ removed spurious detections. Tracks were made using Trackpy, using a 5-pixel search radius and up to 25 frames ahead. For each track, the square of the displacement of the tracked particle were calculated for all pairs of frames and the time difference. To calculate the MSD, the square displacements were binned with multiples of 5 seconds and the MSD were calculated in each bin.

The diffusion coefficient *D* was calculated by fitting the first *N* points in the MSD(Δ*t*) trace with MSD(Δ*t*) = 6DΔ*t* + 6σ^2^− 2*t*_*E*_*D*, where σ is the localization error and *t*_*E*_ is the exposure time ^55^. *N* was determined iteratively as 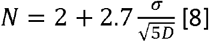,resulting in *D*_U2OS_= 71±5 nm^2^ with *N*= 9 and *D*_yeast_= 239±11 nm^2^/s with *N =*12.

### Focus feedback

A 300 mm cylindrical lens (Thorlabs LJ1558RM-A) was placed in a holder (Zeiss 000000-1772-065) mounted between the Duolink splitter and the front camera (imaging the transcription site or DNA label) resulting in elliptical elongation of any spot within 300 nm of the focal plane ^25,56^. In an identical holder between the Duolink splitter and the second camera (imaging transcription factors), a flat piece of glass as thick as the cylindrical lens was placed to keep the optical path length the same between both channels.

An empirical relation between point spread function (psf) width and distance to the focal plane (z) ^57^ was used to express the ellipticity as a function of z:

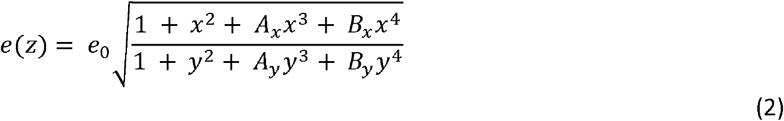

Where 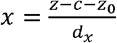 and 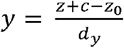. The parameters *e*_0_, *z*_0_, *c, A*_*x*_, *B*_*x*_, *d*_*x*_, *A*_*y*_, *B*_*y*_, and *d*_*y*_ together with *θ*, which gives the angle between a vertically elongated elliptical psf and the vertical of the image, were calibrated using a 100 nm interval z-stack through 200 nm beads (TetraSpeck T7280) mounted in 1% agarose at a concentration of 2.3×10^9^ beads/ml on a Zeiss Coverslip HI coverslip. Each bead in the z-stack was fitted in each plane with an elliptical integrated gaussian psf profile:

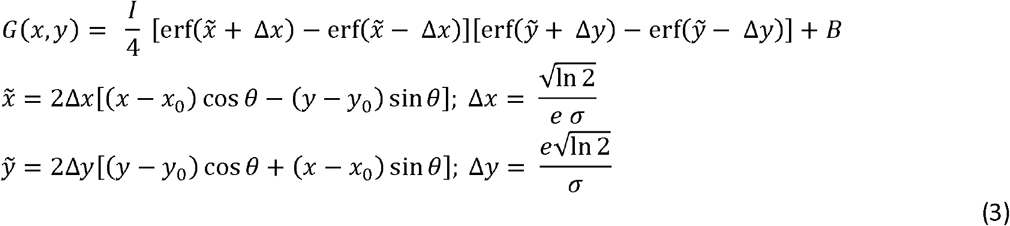

where (*x*_0_, *y*_0_) is the center position of the particle, I its intensity, B the background intensity and e the ellipticity.

After filtering out bad fits, the remainder immediately gives the average *θ* of -0.03. Subsequently, the beads were fitted again with *θ* fixed at -0.03. The ellipticity of each bead was then fitted individually with equation 2 and all beads were aligned such that e(*z*=0) = 1 for each bead. Finally, the data of all beads combined was fitted with equation 2 to yield *e*_0_ = 1.11, z_0_ = -0.12, c = 0.16, *A*_*x*_ = 1.26, *B*_*x*_ = 0.48, *d*_*x*_ = 0.41, *A*_*y*_ = 3.74, *B*_*y*_ = 21.42, *d*_*y*_ = 0.97.

Equation 2 and the calibration were used in a custom-made Python script (10.5281/zenodo.7858759) to correct the focus of the microscope between the acquisition of subsequent frames. Equation 3 was fitted at the location of the brightest pixel in a 48×48 pixel^2^ region of interest centered on the transcription site at the beginning of the experiment. The ellipticity found in this way was used to find z, and the microscope was re-focused by means of a piezo (Zeiss Scanning Stage Piezo 130×100). In experiments with YTL639, the PP7-*GAL10*/PCP-GFPEnvy spot was used for feedback, in experiments with YTL431, the PP7-*RNR2*/PCP-GFPEnvy was used, and for YTL1649 the tetO/tetR-tdTomato DNA label was used.

### Focus feedback performance

The range and error of our focus feedback system were determined by comparing the real position with the position as determined by the focus feedback system (Figure 2C) in a z-stack (101 slices, 100 nm interval) of the sample of TetraSpeck beads described above. The feedback error was defined as the difference between those positions. Its dependence on the distance to the focus can be described as:

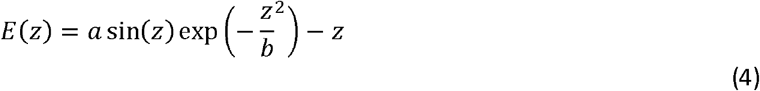

The range of the focus feedback system was defined as two thirds of the distance from *z*=0 to either peak (below to the left and above to the right of *z*=0), as the curve was visibly flat in this range upon inspection of curves with different parameters *a* and *b*. The standard deviation of the error was then calculated over all localizations in this range. Fitting the errors of localizations of beads resulted in a range of 546 nm and a standard deviation of the error in this range of 62 nm.

### Calculating in focus probabilities

The probabilities that a particle is in focus every frame up to some amount of time *T* without (*P*_0_) and with focus feedback (*P*_1_) was calculated assuming diffusion of the particle:

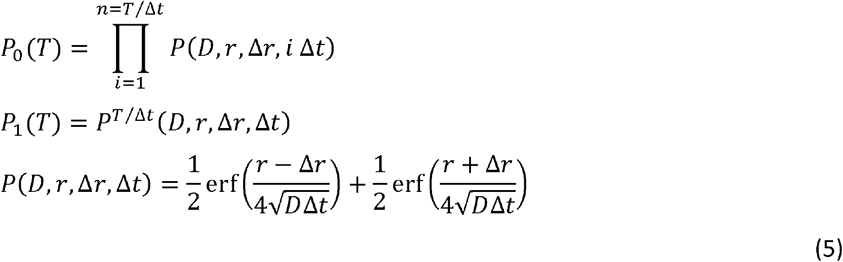

Here D is the diffusion constant; D=239 nm^2^/s and Δ*r* and the range (546 nm) and the error on the feedback (62 nm).

### Channel registration

The cylindrical lens was compressing the image in *x*-direction. To be able to correct for this, for chromatic aberrations and for alignment differences between the two cameras, every day a few z-stacks (101 x 100 nm) of the sample of TetraSpeck beads were acquired as described above. Powers of 0.02%, 0.02% and 0.025% for 561, 488 and 642 nm lasers were used, respectively, and a 33 ms exposure time. An affine transformation between maximum intensity projections of pairs of channels was determined using Simple-Elastix ^58^. In the case of two-channel experiments, the Gal4 channel was not distorted by a cylindrical lens and thus this channel was set to define the fixed grid to which the other channel is to be transformed. In the case of three-channel experiments, all channels were distorted by the cylindrical lens. The distortion by the cylindrical lens was determined once as an 86% scaling in *x* direction. After determining affine transformations of the *GAL10* and Gal4 channels with respect to the DNA label channel, the 86% scaling in *x* direction was applied to all channels. The resulting affine transformations were not applied to the data but saved for use during spot detection (see localization and tracking).

## Statistical analysis

Microscopy data was analyzed using a custom script written in Python (10.5281/zenodo.7858704). The processed source data is available at 10.5281/zenodo.7858747.

### Localization and tracking

In each experiment a mask was drawn for the cell in the center of the image using Otsu thresholding and water shedding on the *GAL10* channel, masks were then transformed to other channels using the affine transformation (see channel registration). The possible spot (DNA label, *GAL10*/*RNR2* TS or Gal4) locations in this cell were determined in each frame using a Laplacian of Gaussian filter with sigma’s 1.91, 1.67 and 2.18 pixels for DNA label, *GAL10*/*RNR2* TS and Gal4, respectively. These detections were then localized using the L-BFGS-B fit algorithm on equation 3 with an additional tilted background:

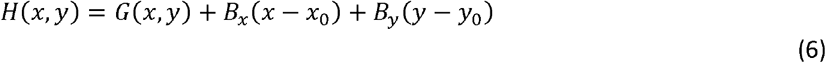

Theta was kept fixed at -0.03 and for channels without cylindrical lens (Gal4 in YTL639 and YTL431 experiments) the ellipticity was fixed at 1. Subsequently, R^2^ and fit confidence intervals were used to remove spurious detections. Positions of the localizations were corrected using the affine transformations (see channel registration).

The same bead stacks used for channel registration were used for correction of spot intensities. Measured integrated intensities of spots decrease with the distance to the focal plane. This dependence was modeled with a Gaussian, which was used to fit the intensity versus distance to focus of all beads in the bead stacks.

The resulting models (one for each day of data acquisition) were used to correct measured intensities back to the in-focus level. Since the distance to focus could only be determined for channels with a cylindrical lens, the correction was only performed on those channels.

Linking of particles in subsequent frames was done using Trackpy (10.5281/zenodo.4682814) searching within 5 pixels and up to 10 frames ahead, for the DNA channel if present and for the *GAL10*/*RNR2* TS channel otherwise. Especially when tracking *GAL10*/*RNR2* TSs, there are frames without a localization of the TS (because there is no burst at the time). To fill the gaps between linked localizations, its position was interpolated in the frame from the previous and next 5 localizations. First it was checked if an unlinked particle or track was present within 5 pixels and if so, it was linked. If not, an attempt was made to fit a new localization within 5 pixels and linked it if successful. If these two failed, a fixed position was fit and linked given by the interpolation.

In experiments with a DNA label, the position of the DNA label was used to find the TS within 3 pixels. In frames where the TS is missing, it was checked if an unlinked particle or track is present within 3 pixels of the DNA label and if so, it was linked. If not, an attempt was made to fit a new localization within 3 pixels and linked it if successful. If these two failed, a fixed position given by the DNA label was fit and linked.

The position of the nearest TF was found in the same way within 3 pixels of the DNA label if present and within 3 pixels of the TS otherwise. 3 pixels, which equates to 300 nm, was shown to be the range in which the majority of close-proximity TF’s can be found ^29^.

To estimate the background intensity surrounding the spot, equation 6 was fit in the four cardinal directions around each localization at 5 pixels distance. In case more than one TS or DNA label track was found, for example due to a linked track of spurious localizations, the brightest track was kept. The results were plotted on crops of the raw data which were used to determine the appropriate range of frames to use for later analysis. For each experiment, the first 30 frames were excluded. In these 30 frames high (auto)fluorescence was visible in the 640 nm (TF) channel, skewing spot intensities. In some experiments we excluded several frames at the end because no TF was visible anymore because of photobleaching.

### Burst and TF binding binarization

The ON and OFF periods of transcription and the periods with and without detected TF binding were modeled by a hidden Markov model (HMM). An initial probability distribution for transcriptional bursting is given by a normal distribution centered on 500 counts and with a 300-count standard deviation. An initial probability distribution for the TF bound state is given by a normal distribution centered on 100 counts and with a 250-count standard deviation. These probability distributions are then optimized by the Baum-Welch algorithm for each experiment individually and subsequently used in the Viterbi algorithm to predict burst ON and OFF states and TF bound and unbound/undetected states.

### TF intensities

The intensities of the detected Gal4 binding events provide insight into the number of detected Gal4 molecules using dense labeling (in YTL639) and sparse labeling (in YTL1649) (Figure S1E). The TF tracks (see Localization and tracking) were filtered on periods where the TF was bound (see Burst and TF binding binarization). The intensities of the TFs in those frames were then divided by 0.4 (YTL639) or 1 (YTL1649) to account for the difference in illumination power.

### Burst and TF binding alignment

To align on the starts of each burst, each intensity trace was copied n times according to the number of bursts in the trace. On each of these copies, *t*=0 was defined at subsequent bursts from the original trace. This alignment was done for all traces in an experiment. After aligning (by redefining *t=*0), an average intensity trace was made by calculating a mean intensity and standard error of the mean for each time point according to their newly defined time with respect to the start of a burst. A similar approach was used to align to the ends of bursts and to the starts and ends of TF binding events. Errors were determined with bootstrapping with 10.000 repeats ^59^.

### Determining Gal4 transcription activation efficiency

The fraction of Gal4 binding events resulting in transcription activation (Figure 5A) was calculated by aligning TF intensity traces at the starts of TF binding events. A Savitzky-Golay filter of order 3 and with a window of 5 frames was used to calculate the first derivative of the TS intensity. The fraction (*R*) of TS intensities that was increasing at each aligned time frame was determined. This fraction was compared with the fraction of increasing TS intensities at random moments during bursts (*B*), which was always very close to 0.5. The correction was done as *F* = (*R−B*)/(1−*B*).

To determine the initiation rate, the PP7-*GAL10* signal was converted to the number of nascent transcripts by comparison of the distribution of live-cell TS intensities during bursts, as found by HMM binarization, to number of mRNA distributions, determined by smFISH (https://osf.io/d5nvj/)^26^, (Figure S5A). An interpolated cumulative distribution function (cdf) of TS intensities was fit to an interpolated cdf of number of mRNA from smFISH, by minimizing their sum of square differences, multiplied by the scaling factor and an added offset, resulting in a factor of 1.06 ± 0.28×10^−2^ counts/mRNA and an offset of 9.29 ± 0.94 mRNA. The initiation rate (Figure 5B) was then calculated by aligning the normalized TS intensity traces at the starts of bursts. A Savitzky-Golay filter of order 3 and with a window of 5 frames was used to calculate the first derivative of the number of mRNAs. This derivative was then averaged. Errors were determined with bootstrapping with 10.000 repeats ^59^.

### Survival time calculation

The durations of the burst ON and OFF times were determined, as well as TF binding durations and time between TF binding events from the binarized traces. The distributions of those time were plotted as survival distributions. Simply plotting the number of surviving events at each duration would exclude events bordering the start or end of the trace because the exact duration of these border-events is unknown. To circumvent this, the number of surviving events (including events bordering starts and ends of traces) was divided by the number of events which have at least the required duration. The resulting curve was fit by both one- and two fraction exponential distributions:

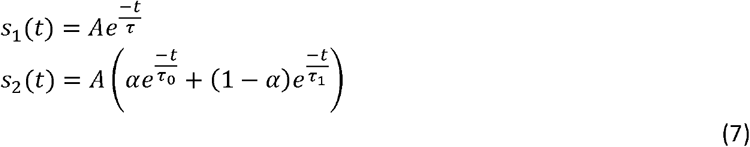

Which function is describing the curve best is determined using Bayes Information Criterion.

### Determining TF occupancy and association/dissociation frequencies

Each frame in the intensity traces was classified into five periods (Figure 6A) according to the following rules:

- Before a burst: at least 25s after the previous burst until 10 s before the start of a burst
- At the start of a burst: between 10 s before the start until 10 s after the start of a burst and at least 25 s after the end of the previous burst.
- During a burst: between 10 s after the start of a burst until 10 s before the end of a burst.
- At the end of a burst: between 10 s before until 10 s after the end of a burst and at least 25 s before the start of the next burst.
- After a burst: At least 10 s after the end of a burst and at least 25 s before the start of the next burst. For all frames in these five periods, the TF occupancy was determined by dividing the number of frames with a TF bound by the total number of frames in each period. To obtain association and dissociation frequencies, the number of TF association or dissociation events were divided by the number of time frames in each period, respectively. Association and dissociation frequencies were calculated for all binding TF events and for TF binding events longer than 20 s to enrich for *GAL10*-specific binding events. Standard errors of the means of all these numbers as well as the p-values shown were calculated by bootstrapping over 100.000 repeats ^59^. Multiple testing correction was performed using the Bonferroni method.

### Overlap between bursts and binding events

The fraction of bursts overlapping with the longest binding event was determined as follows. First, for each burst the TF binding event overlapping the greatest number of frames with the burst was found. Second, the overlap divided by the burst duration: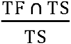. Different sets of bursts were selected consisting only of bursts with lengths of at least 0, 10, 20, 40, 55, 75 and 100 s.

The fraction of TF binding overlapping with a burst was calculated analogously. The burst overlapping the greatest number of frames with the TF binding event was found and the overlap was divided by the burst duration: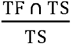. Different sets of binding events were selected consisting only of events with durations of at least 0, 10, 20, 40, 55, 75 and 100 s.

### Number of binding events per burst

The number of binding events per burst (Figure S7) was obtained by counting the number of binding events which overlap with the burst at least one frame. The bursts found in YTL1649 were divided into two equal groups, with bursts durations of either less (143 bursts) or more (140 bursts) than 50 s. The same was done for the control YTL431, splitting into two groups of 56 and 81 bursts respectively. To estimate whether distributions were significantly different, *p*-values were determined between YTL1649 and YTL431 distributions for both bursts duration regimes by bootstrapping with 10.000 repeats ^59^.

### Correlation functions

The average cross-correlation function (ccf) is a measure for correlation between two time series when one of the time series has been shifted by a time delay *τ*. Analyzed frames were defined with a mask m(t), which is 1 when the frame at time t is included and 0 otherwise. Defining the TS intensity traces as *a*_i_ and the TF intensity traces as *b*_i_, the ccf’s were calculated for each trace i as follows:

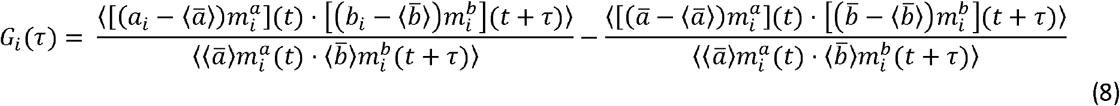

Here, averages were denoted over t with ⟨ ⟩ and averages over all traces (*i*) with ⍰. The average ccf ⍰ (*τ*) over the ccfs were calculated as 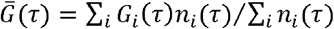, where *n_i_* (τ) is the number of possible pairs of timepoints in the *i*^th^ TS and TF intensity traces. The correlation theorem and fast Fourier transforms were used to speed up the calculation of *G_i_*(*τ*). The standard errors of the mean were determined by bootstrapping with 10.000 repeats ^59^.

## Notes

### Competing Interest Statement

The authors have declared no competing interest.

## References

1. Mazzocca, M., Fillot, T., Loffreda, A., Gnani, D., and Mazza, D. (2021). The needle and the haystack: single molecule tracking to probe the transcription factor search in eukaryotes. Biochem. Soc. Trans. 49, 1121–1132. 10.1042/BST20200709.

2. Jana, T., Brodsky, S., and Barkai, N. (2021). Speed-Specificity Trade-Offs in the Transcription Factors Search for Their Genomic Binding Sites. Trends Genet. TIG 37, 421–432. 10.1016/j.tig.2020.12.001.

3. Ferrie, J.J., Karr, J.P., Tjian, R., and Darzacq, X. (2022). “Structure”-function relationships in eukaryotic transcription factors: The role of intrinsically disordered regions in gene regulation. Mol. Cell 82, 3970–3984. 10.1016/j.molcel.2022.09.021.

4. Lu, F., and Lionnet, T. (2021). Transcription Factor Dynamics. Cold Spring Harb. Perspect. Biol. 13, a040949. 10.1101/cshperspect.a040949.

5. Rodriguez, J., and Larson, D.R. (2020). Transcription in Living Cells: Molecular Mechanisms of Bursting. Annu. Rev. Biochem. 89, 189–212. 10.1146/annurev-biochem-011520-105250.

6. Brouwer, I., and Lenstra, T.L. (2019). Visualizing transcription: key to understanding gene expression dynamics. Curr. Opin. Chem. Biol. 51, 122–129. 10.1016/j.cbpa.2019.05.031.

7. Larson, D.R., Fritzsch, C., Sun, L., Meng, X., Lawrence, D.S., and Singer, R.H. (2013). Direct observation of frequency modulated transcription in single cells using light activation. eLife 2, e00750. 10.7554/eLife.00750.

8. Loffreda, A., Jacchetti, E., Antunes, S., Rainone, P., Daniele, T., Morisaki, T., Bianchi, M.E., Tacchetti, C., and Mazza, D. (2017). Live-cell p53 single-molecule binding is modulated by C-terminal acetylation and correlates with transcriptional activity. Nat. Commun. 8, 313. 10.1038/s41467-017-00398-7.

9. Rullan, M., Benzinger, D., Schmidt, G.W., Milias-Argeitis, A., and Khammash, M. (2018). An Optogenetic Platform for Real-Time, Single-Cell Interrogation of Stochastic Transcriptional Regulation. Mol. Cell 70, 745–756.e6. 10.1016/j.molcel.2018.04.012.

10. Senecal, A., Munsky, B., Proux, F., Ly, N., Braye, F.E., Zimmer, C., Mueller, F., and Darzacq, X. (2014). Transcription factors modulate c-Fos transcriptional bursts. Cell Rep. 8, 75–83. 10.1016/j.celrep.2014.05.053.

11. de Jonge, W.J., Patel, H.P., Meeussen, J.V.W., and Lenstra, T.L. (2022). Following the tracks: How transcription factor binding dynamics control transcription. Biophys. J. 121, 1583–1592. 10.1016/j.bpj.2022.03.026.

12. Lammers, N.C., Kim, Y.J., Zhao, J., and Garcia, H.G. (2020). A matter of time: Using dynamics and theory to uncover mechanisms of transcriptional bursting. Curr. Opin. Cell Biol. 67, 147–157. 10.1016/j.ceb.2020.08.001.

13. Mazzocca, M., Colombo, E., Callegari, A., and Mazza, D. (2021). Transcription factor binding kinetics and transcriptional bursting: What do we really know? Curr. Opin. Struct. Biol. 71, 239–248. 10.1016/j.sbi.2021.08.002.

14. Boija, A., Klein, I.A., Sabari, B.R., Dall’Agnese, A., Coffey, E.L., Zamudio, A.V., Li, C.H., Shrinivas, K., Manteiga, J.C., Hannett, N.M., et al. (2018). Transcription Factors Activate Genes through the Phase-Separation Capacity of Their Activation Domains. Cell 175, 1842–1855.e16. 10.1016/j.cell.2018.10.042.

15. Chong, S., Dugast-Darzacq, C., Liu, Z., Dong, P., Dailey, G.M., Cattoglio, C., Heckert, A., Banala, S., Lavis, L., Darzacq, X., et al. (2018). Imaging dynamic and selective low-complexity domain interactions that control gene transcription. Science 361. 10.1126/science.aar2555.

16. Popp, A.P., Hettich, J., and Gebhardt, J.C.M. (2021). Altering transcription factor binding reveals comprehensive transcriptional kinetics of a basic gene. Nucleic Acids Res. 49, 6249–6266. 10.1093/nar/gkab443.

17. Stavreva, D.A., Garcia, D.A., Fettweis, G., Gudla, P.R., Zaki, G.F., Soni, V., McGowan, A., Williams, G., Huynh, A., Palangat, M., et al. (2019). Transcriptional Bursting and Co-bursting Regulation by Steroid Hormone Release Pattern and Transcription Factor Mobility. Mol. Cell 75, 1161–1177.e11. 10.1016/j.molcel.2019.06.042.

18. Donovan, B.T., Huynh, A., Ball, D.A., Patel, H.P., Poirier, M.G., Larson, D.R., Ferguson, M.L., and Lenstra, T.L. (2019). Live-cell imaging reveals the interplay between transcription factors, nucleosomes, and bursting. EMBO J. 38. 10.15252/embj.2018100809.

19. Garcia, D.A., Fettweis, G., Presman, D.M., Paakinaho, V., Jarzynski, C., Upadhyaya, A., and Hager, G.L. (2021). Power-law behavior of transcription factor dynamics at the single-molecule level implies a continuum affinity model. Nucleic Acids Res. 49, 6605–6620. 10.1093/nar/gkab072.

20. Bertrand, E., Chartrand, P., Schaefer, M., Shenoy, S.M., Singer, R.H., and Long, R.M. (1998). Localization of ASH1 mRNA particles in living yeast. Mol. Cell 2, 437–445.

21. Larson, D.R., Zenklusen, D., Wu, B., Chao, J.A., and Singer, R.H. (2011). Real-time observation of transcription initiation and elongation on an endogenous yeast gene. Science 332, 475–478. 10.1126/science.1202142.

22. Rhee, H.S., and Pugh, B.F. (2011). Comprehensive genome-wide protein-DNA interactions detected at single-nucleotide resolution. Cell 147, 1408–1419. 10.1016/j.cell.2011.11.013.

23. Grimm, J.B., English, B.P., Chen, J., Slaughter, J.P., Zhang, Z., Revyakin, A., Patel, R., Macklin, J.J., Normanno, D., Singer, R.H., et al. (2015). A general method to improve fluorophores for live-cell and single-molecule microscopy. Nat. Methods 12, 244–250, 3 p following 250. 10.1038/nmeth.3256.

24. Tokunaga, M., Imamoto, N., and Sakata-Sogawa, K. (2008). Highly inclined thin illumination enables clear single-molecule imaging in cells. Nat. Methods 5, 159–161. 10.1038/nmeth1171.

25. Kao, H.P., and Verkman, A.S. (1994). Tracking of single fluorescent particles in three dimensions: use of cylindrical optics to encode particle position. Biophys. J. 67, 1291–1300. 10.1016/S0006-3495(94)80601-0.

26. Fu, X., Patel, H.P., Coppola, S., Xu, L., Cao, Z., Lenstra, T.L., and Grima, R. (2022). Quantifying how post-transcriptional noise and gene copy number variation bias transcriptional parameter inference from mRNA distributions. eLife 11, e82493. 10.7554/eLife.82493.

27. Mazza, D., Abernathy, A., Golob, N., Morisaki, T., and McNally, J.G. (2012). A benchmark for chromatin binding measurements in live cells. Nucleic Acids Res. 40, e119. 10.1093/nar/gks701.

28. Ball, D.A., Mehta, G.D., Salomon-Kent, R., Mazza, D., Morisaki, T., Mueller, F., McNally, J.G., and Karpova, T.S. (2016). Single molecule tracking of Ace1p in Saccharomyces cerevisiae defines a characteristic residence time for non-specific interactions of transcription factors with chromatin. Nucleic Acids Res. 10.1093/nar/gkw744.

29. Meeussen, J.V.W., Pomp, W., Brouwer, I., de Jonge, W.J., Patel, H.P., and Lenstra, T.L. (2023). Transcription factor clusters enable target search but do not contribute to target gene activation. Nucleic Acids Res., gkad227. 10.1093/nar/gkad227.

30. Dovrat, D., Dahan, D., Sherman, S., Tsirkas, I., Elia, N., and Aharoni, A. (2018). A Live-Cell Imaging Approach for Measuring DNA Replication Rates. Cell Rep. 24, 252–258. 10.1016/j.celrep.2018.06.018.

31. Lickwar, C.R., Mueller, F., Hanlon, S.E., McNally, J.G., and Lieb, J.D. (2012). Genome-wide protein-DNA binding dynamics suggest a molecular clutch for transcription factor function. Nature 484, 251–255. 10.1038/nature10985.

32. Kang, T., Martins, T., and Sadowski, I. (1993). Wild type GAL4 binds cooperatively to the GAL1-10 UASG in vitro. J. Biol. Chem. 268, 9629–9635.

33. Giniger, E., and Ptashne, M. (1988). Cooperative DNA binding of the yeast transcriptional activator GAL4. Proc. Natl. Acad. Sci. U. S. A. 85, 382–386.

34. Adams, C.C., and Workman, J.L. (1995). Binding of disparate transcriptional activators to nucleosomal DNA is inherently cooperative. Mol. Cell. Biol. 15, 1405–1421. 10.1128/MCB.15.3.1405.

35. Ibarra, I.L., Hollmann, N.M., Klaus, B., Augsten, S., Velten, B., Hennig, J., and Zaugg, J.B. (2020). Mechanistic insights into transcription factor cooperativity and its impact on protein-phenotype interactions. Nat. Commun. 11, 124. 10.1038/s41467-019-13888-7.

36. Brouwer, I., Kerklingh, E., van Leeuwen, F., and Lenstra, T.L. (2023). Dynamic epistasis analysis reveals how chromatin remodeling regulates transcriptional bursting. Nat. Struct. Mol. Biol. 10.1038/s41594-023-00981-1.

37. Jolma, A., Yin, Y., Nitta, K.R., Dave, K., Popov, A., Taipale, M., Enge, M., Kivioja, T., Morgunova, E., and Taipale, J. (2015). DNA-dependent formation of transcription factor pairs alters their binding specificity. Nature 527, 384–388. 10.1038/nature15518.

38. Sönmezer, C., Kleinendorst, R., Imanci, D., Barzaghi, G., Villacorta, L., Schübeler, D., Benes, V., Molina, N., and Krebs, A.R. (2021). Molecular Co-occupancy Identifies Transcription Factor Binding Cooperativity In Vivo. Mol. Cell 81, 255–267.e6. 10.1016/j.molcel.2020.11.015.

39. Rao, S., Ahmad, K., and Ramachandran, S. (2021). Cooperative binding between distant transcription factors is a hallmark of active enhancers. Mol. Cell 81, 1651–1665.e4. 10.1016/j.molcel.2021.02.014.

40. Iyer, V., and Struhl, K. (1996). Absolute mRNA levels and transcriptional initiation rates in Saccharomyces cerevisiae. Proc. Natl. Acad. Sci. U. S. A. 93, 5208–5212.

41. Pelechano, V., Chávez, S., and Pérez-Ortín, J.E. (2010). A complete set of nascent transcription rates for yeast genes. PloS One 5, e15442. 10.1371/journal.pone.0015442.

42. Nguyen, V.Q., Ranjan, A., Liu, S., Tang, X., Ling, Y.H., Wisniewski, J., Mizuguchi, G., Li, K.Y., Jou, V., Zheng, Q., et al. (2021). Spatiotemporal coordination of transcription preinitiation complex assembly in live cells. Mol. Cell 81, 3560–3575.e6. 10.1016/j.molcel.2021.07.022.

43. Xiong, L., Zeng, Y., Tang, R.-Q., Alper, H.S., Bai, F.-W., and Zhao, X.-Q. (2018). Condition-specific promoter activities in Saccharomyces cerevisiae. Microb. Cell Factories 17, 58. 10.1186/s12934-018-0899-6.

44. Darzacq, X., Shav-Tal, Y., de Turris, V., Brody, Y., Shenoy, S.M., Phair, R.D., and Singer, R.H. (2007). In vivo dynamics of RNA polymerase II transcription. Nat. Struct. Mol. Biol. 14, 796–806. 10.1038/nsmb1280.

45. Steurer, B., Janssens, R.C., Geverts, B., Geijer, M.E., Wienholz, F., Theil, A.F., Chang, J., Dealy, S., Pothof, J., van Cappellen, W.A., et al. (2018). Live-cell analysis of endogenous GFP-RPB1 uncovers rapid turnover of initiating and promoter-paused RNA Polymerase II. Proc. Natl. Acad. Sci. U. S. A. 115, E4368–E4376. 10.1073/pnas.1717920115.

46. Pimmett, V.L., Dejean, M., Fernandez, C., Trullo, A., Bertrand, E., Radulescu, O., and Lagha, M. (2021). Quantitative imaging of transcription in living Drosophila embryos reveals the impact of core promoter motifs on promoter state dynamics. Nat. Commun. 12, 4504. 10.1038/s41467-021-24461-6.

47. Zoller, B., Little, S.C., and Gregor, T. (2018). Diverse Spatial Expression Patterns Emerge from Unified Kinetics of Transcriptional Bursting. Cell 175, 835–847.e25. 10.1016/j.cell.2018.09.056.

48. Rodriguez, J., Ren, G., Day, C.R., Zhao, K., Chow, C.C., and Larson, D.R. (2019). Intrinsic Dynamics of a Human Gene Reveal the Basis of Expression Heterogeneity. Cell 176, 213–226.e18. 10.1016/j.cell.2018.11.026.

49. Lenstra, T.L., Coulon, A., Chow, C.C., and Larson, D.R. (2015). Single-Molecule Imaging Reveals a Switch between Spurious and Functional ncRNA Transcription. Mol. Cell 60, 597–610. 10.1016/j.molcel.2015.09.028.

50. Patel, H.P., Coppola, S., Pomp, W., Brouwer, I., and Lenstra, T.L. (2022). DNA supercoiling restricts the transcriptional bursting of neighboring eukaryotic genes. 2022.03.04.482969. 10.1101/2022.03.04.482969.

51. Laughery, M.F., Hunter, T., Brown, A., Hoopes, J., Ostbye, T., Shumaker, T., and Wyrick, J.J. (2015). New vectors for simple and streamlined CRISPR-Cas9 genome editing in Saccharomyces cerevisiae. Yeast Chichester Engl. 32, 711–720. 10.1002/yea.3098.

52. Brouwer, I., Patel, H.P., Meeussen, J.V.W., Pomp, W., and Lenstra, T.L. (2020). Single-Molecule Fluorescence Imaging in Living Saccharomyces cerevisiae Cells. STAR Protoc. 1, 100142. 10.1016/j.xpro.2020.100142.

53. Grimm, J.B., Xie, L., Casler, J.C., Patel, R., Tkachuk, A.N., Falco, N., Choi, H., Lippincott-Schwartz, J., Brown, T.A., Glick, B.S., et al. (2021). A General Method to Improve Fluorophores Using Deuterated Auxochromes. JACS Au 1, 690–696. 10.1021/jacsau.1c00006.

54. Palangat, M., and Larson, D.R. (2016). Single-gene dual-color reporter cell line to analyze RNA synthesis in vivo. Methods San Diego Calif 103, 77–85. 10.1016/j.ymeth.2016.04.009.

55. Michalet, X. (2010). Mean square displacement analysis of single-particle trajectories with localization error: Brownian motion in an isotropic medium. Phys. Rev. E 82, 041914. 10.1103/PhysRevE.82.041914.

56. Holtzer, L., Meckel, T., and Schmidt, T. (2007). Nanometric three-dimensional tracking of individual quantum dots in cells. Appl. Phys. Lett. 90, 053902. 10.1063/1.2437066.

57. Huang, B., Wang, W., Bates, M., and Zhuang, X. (2008). Three-dimensional super-resolution imaging by stochastic optical reconstruction microscopy. Science 319, 810–813. 10.1126/science.1153529.

58. Marstal, K., Berendsen, F., Staring, M., and Klein, S. (2016). SimpleElastix: A User-Friendly, Multi-lingual Library for Medical Image Registration. In 2016 IEEE Conference on Computer Vision and Pattern Recognition Workshops (CVPRW), pp. 574–582. 10.1109/CVPRW.2016.78.

59. Efron, Bradley and Tibshirani, Robert J. (1993). Chapter 16: Hypothesis testing with the bootstrap, in An Introduction to the Bootstrap (Springer-Science+Business Media, B.V.).

