## Supplemental figures for "Transcription factor exchange enables prolonged transcriptional bursts"

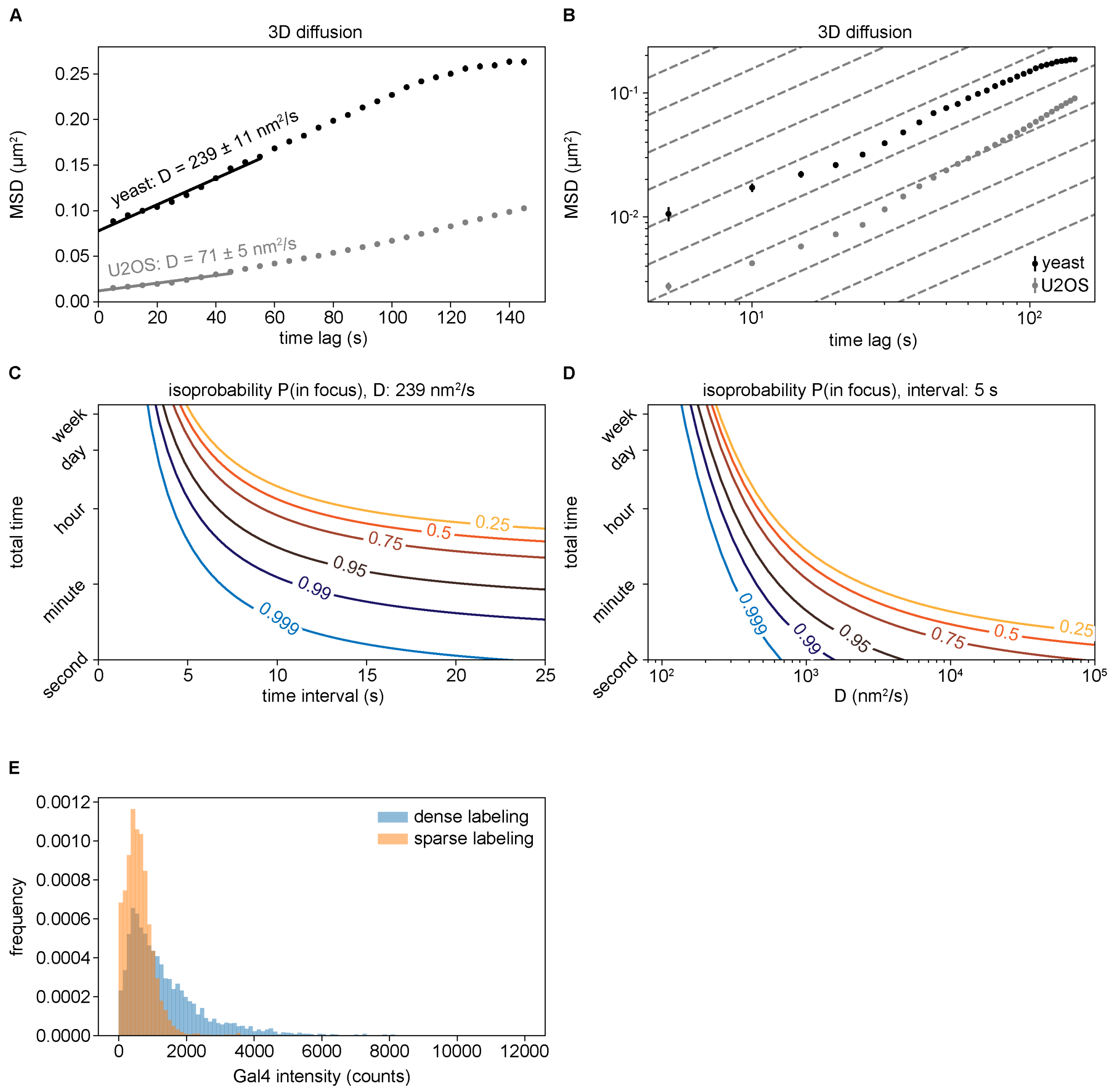


**Figure S1. Real-time tracking using active feedback**

**A.** Mean Squared Displacement plots for a transcription site in budding yeast (black) and U2OS (grey) cells, obtained from 3D imaging. Measured diffusion constants (D) are indicated. The diffusion constant includes any movement of the tracked transcription site with respect to the objective, such as movement and rotation of the nucleus, movement of the cell, the sample and drift in the microscope stage.

**B.** Same as A, plotted with log-log axis, which shows that the slope ($\alpha$) at short timescales is less than 1 (dotted grey lines), in line with subdiffusive movement of chromatin loci within the nucleus. At longer timescales, movement and rotation of the nucleus, movement of the cell, the sample and drift in the microscope stage cause deviation of this slope. The tracking corrects for all these movements regardless of their origin.

**C.** Lines of constant probability that a particle is in focus in all frames as a function of time interval between frames and total acquisition time.

**D.** Lines of constant probability that a particle is in focus in all frames as a function of diffusion constant and total acquisition time.

**E.** Histogram of Gal4 intensity distribution during Gal4 bound periods of the Gal4 tracks, as determined with a HMM. The intensity was adjusted for differences in illumination power used for the dense and sparse labeling experiments (methods). The sparse labeling (orange) showed a unimodal narrow peak, indicating single molecules. The dense labeling (blue) showed similar or higher intensities, indicating detected Gal4 binding events represented single or multiple molecules.

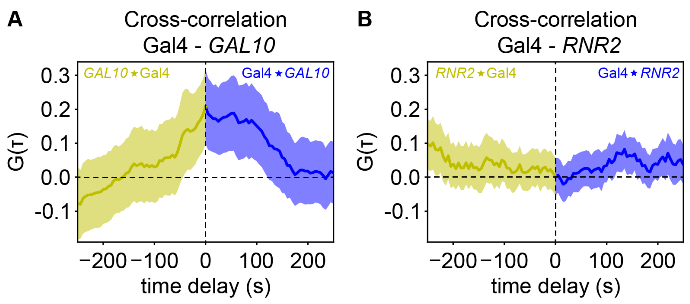


**Figure S2: Gal4 binding and *GAL10* transcription are dynamically coupled.**

**A.** Cross-correlation of Gal4-*GAL10*. Gal4 ⋆ *GAL10* indicates the cross-correlation function of Gal4(t) to *GAL10*(t+τ) (blue) and *GAL10* ⋆ Gal4 indicates the cross-correlation function of Gal4(t-τ) to *GAL10*(t) (yellow). *n* = 46 cells.

**B.** Cross-correlation of Gal4-*RNR2*. Gal4 ⋆ *RNR2* indicates the cross-correlation function of Gal4(t) to *RNR2*(t+τ) (blue) and *RNR2* ⋆ Gal4 indicates the cross-correlation function of Gal4(t-τ) to *RNR2*(t) (yellow). *n* = 34 cells.

Shaded area indicates standard error of the mean.

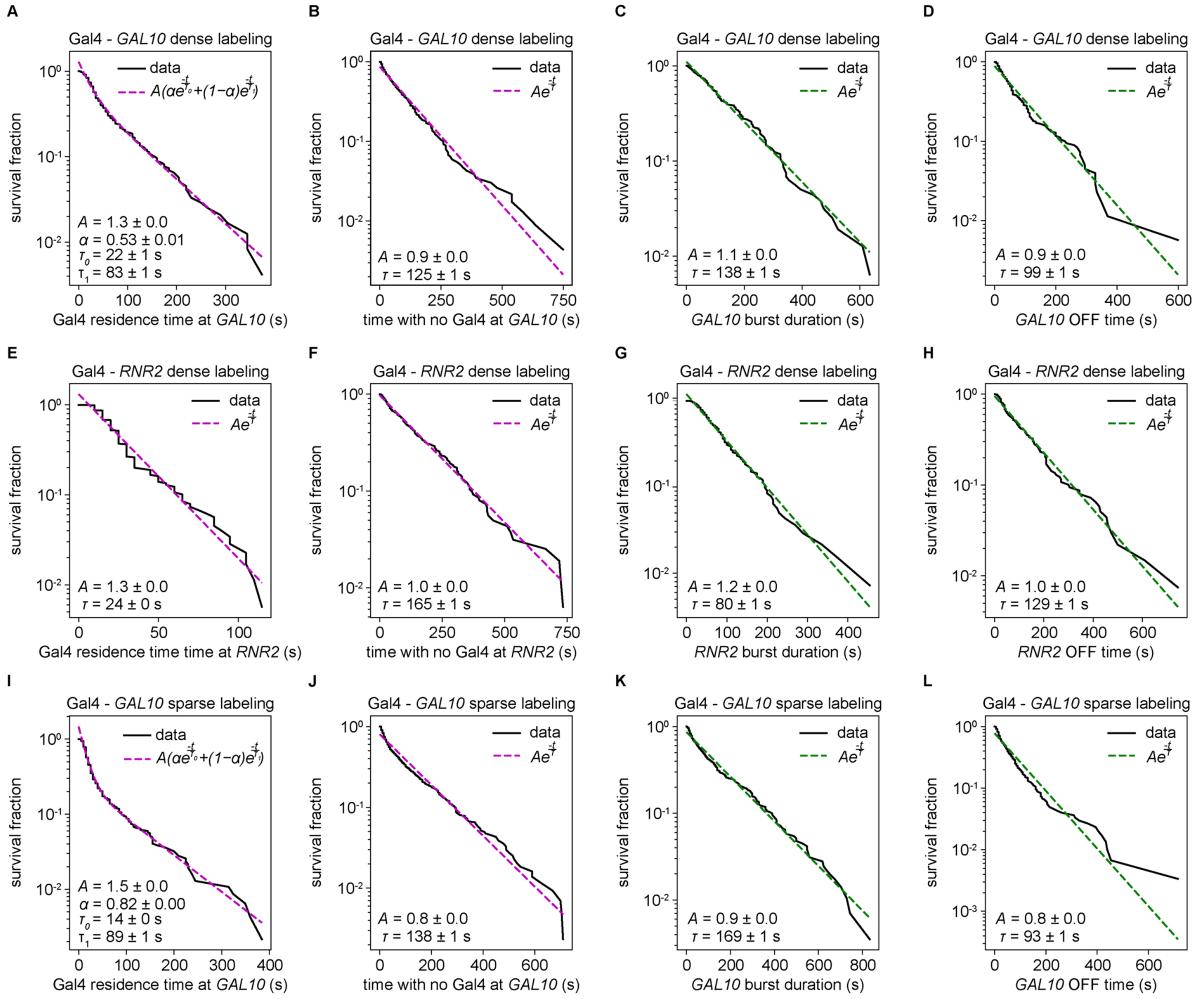


**Figure S3. Survival probability of Gal4 residence times, times with no detected Gal4 binding events and *GAL10*/*RNR2* ON and OFF periods for different experiments.**

**A.** Survival fraction (*s*/*r*) of the residence time of HMM-binarized Gal4 residence times at the *GAL10* locus, using dense Gal4 labeling.

**B.** Survival fraction (*s*/*r*) of the HMM-binarized periods with no detected Gal4 binding events at the *GAL10* locus, which represent periods where Gal4 is either unbound or unlabeled.

**C.** Survival fraction (*s*/*r*) of the HMM-binarized *GAL10* burst duration (ON times).

**D.** Survival fraction (*s*/*r*) of the HMM-binarized *GAL10* OFF periods.

**E-H.** Same as A-D for the *RNR2* gene, using dense Gal4 labeling.

**I-L.** Same as A-D for the *GAL10* gene, using sparse Gal4 labeling.

The probability (*s*) is corrected for the number of traces (*r*) for each residence time. We used Bayes Information Criteria to determine whether the survival distribution was best fit with an exponential or a biexponential distribution. Only the best fit is shown as a dotted line with the indicated parameters. Errors indicate standard errors of the fit. For A-D: $n$ = 46 cells, for E-H: $n$ = 34 cells, for I-L: $n$ = 79 cells

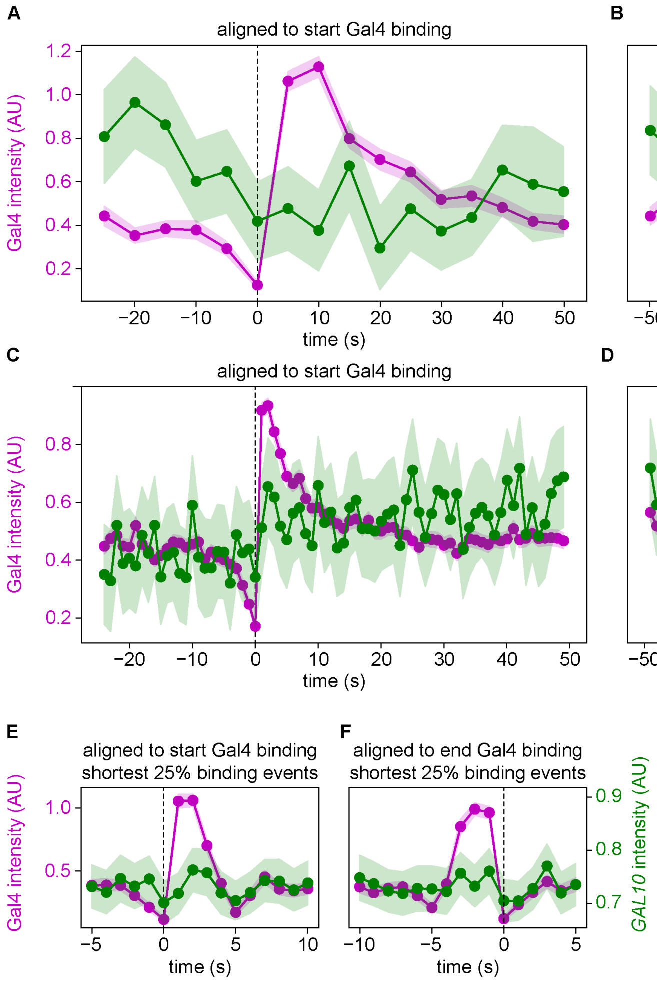


**Figure S4. Transcription factor association and dissociation determine burst start and end, respectively.**

**A.** Average Gal4 (magenta) and *RNR2* TS (green) intensities aligned to the start of Gal4 binding.

**B.** Same as A aligned to the end of Gal4 binding.

**C.** Average Gal4 (magenta) and *GAL10* TS (green) intensities aligned to the start of Gal4 binding after tracking with a 1 second interval.

**D.** Same as F aligned to the end of Gal4 binding.

**E.** Same as C, using only the 25% shortest Gal4 binding events

**F.** Same as E, aligned to the end of Gal4 binding

Shaded areas indicate standard errors of the means. For A-B: $n$ = 34 cells, for C-F: $n$ = 55 cells

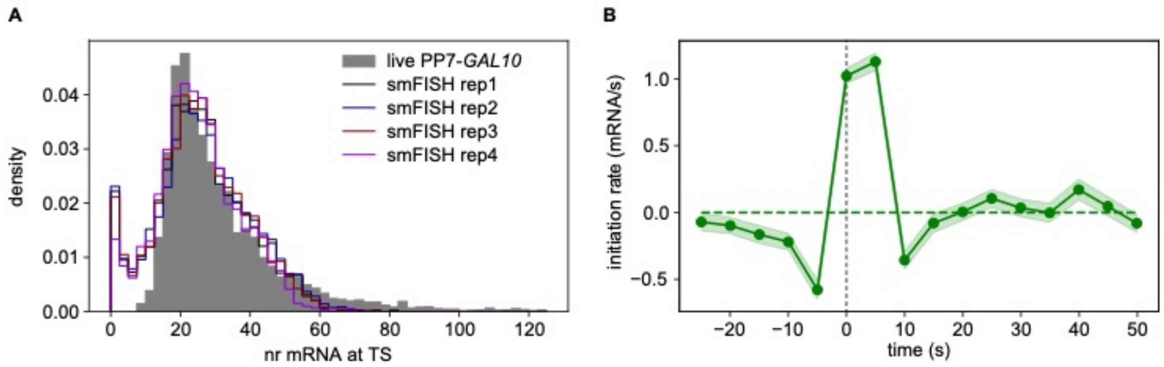


**Figure S5. The TF Gal4 activates transcription efficiently**

**A.** Fitting of live cell PP7-*GAL10* intensity during ON periods (grey) with 4 smFISH datasets[1] hybridized with PP7 probes (colored lines) to convert the measured PP7-*GAL10* intensity to number of RNAs at the TS.

**B.** *GAL10* initiation rate, as measured by the average increase of TS intensity, when tracking was done using a DNA label and *GAL10* TS intensity was measured in a second channel. Data is aligned at the starts of *GAL10* bursts. Shaded area indicates standard error of the mean. *n* = 79 cells

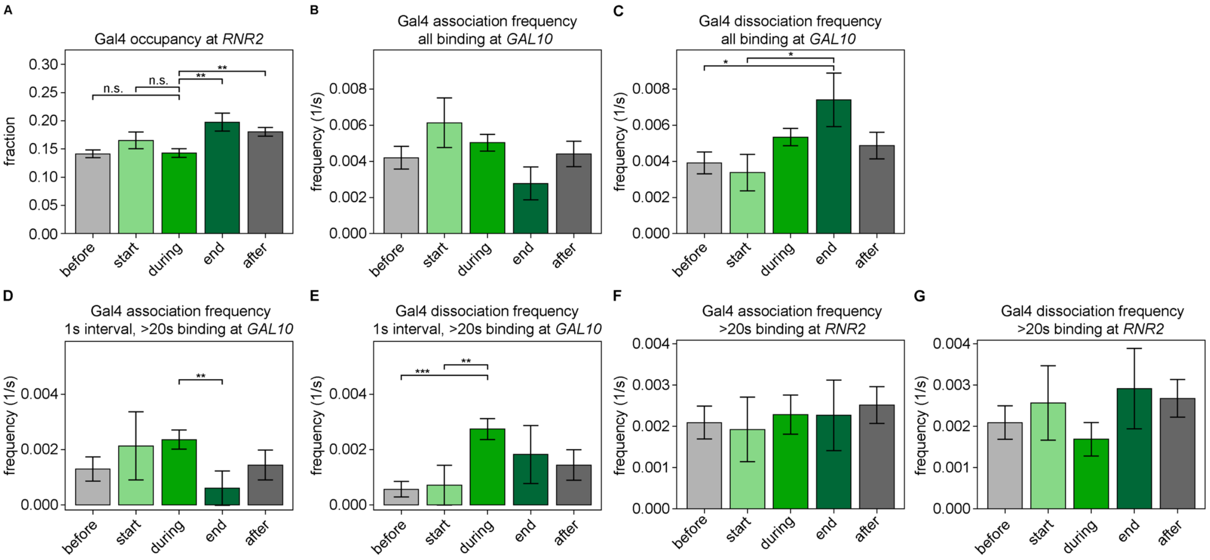


**Figure S6. Gal4 molecules exchange during a transcriptional burst.**

**A.** Gal4 occupancy at *RNR2*, as measured by the fraction of timepoints with bound Gal4, in the time periods indicated in Figure 6A.

**B.** Gal4 association frequency at *GAL10*, as measured by the number of Gal4 association events divided by number of frames, when all Gal4 binding events are considered, in the time periods indicated in Figure 6A.

**C.** Gal4 dissociation frequency at *GAL10*, as measured by the number of Gal4 dissociation events divided by the number of frames, when all Gal4 binding events are considered, in the time periods indicated in Figure 6A.

**D.** Same as B, from tracking with a 1 second interval.

**E.** Same as C, from tracking with a 1 second interval.

**F.** Gal4 association frequency at *RNR2.*

**G.** Gal4 dissociation frequency at *RNR2*.

In D-G only binding events lasting longer than 20 seconds are considered. Error bars indicate standard error of the mean. In A, significance is only calculated for “during at burst” versus all other periods, in B-G, significance was calculated between all time periods, and only significant bars are shown. Significance was determined by bootstrapping with 100.000 repeats, and corrected for multiple testing using the Bonferroni method. * $p < 0.05$, ** $p < 0.005$, *** $p < 0.0005$

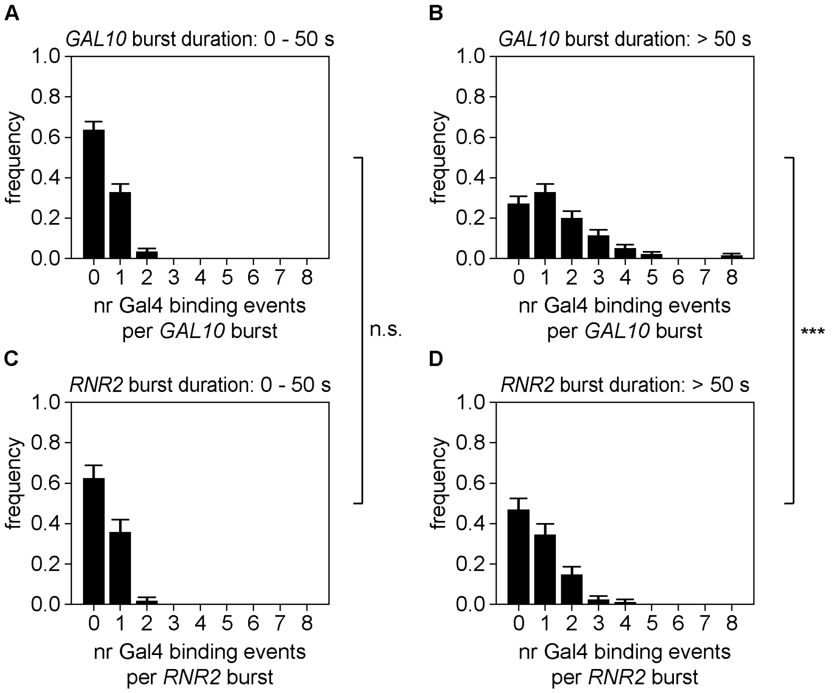


**Figure S7. Longer *GAL10* bursts have more Gal4 binding events**

**A.** Number of Gal4 binding events during short *GAL10* bursts (0 - 50s).

**B.** Number of Gal4 binding events during long *GAL10* bursts (> 50 s).

**C.** Same as A for short *RNR2* bursts.

**D.** Same as B for long *RNR2* bursts.

In A-B Gal4 was sparsely labeled, in C-D Gal4 was densely labeled. Errors indicate standard error of the mean. Significance and errors were determined by bootstrap with 10.000 bootstrap repeats. *** $p < 0.0005$

**Video S1. Simultaneous imaging of Gal4 binding and *GAL10* transcription using focus feedback of the *GAL10* transcription site.**

Example video of a *GAL10* TS, tracked and imaged every 5 s with focus feedback, while simultaneously imaging DNA binding of Gal4-HALO-JF646. In this experiment, Gal4 was labeled densely.
